# Co-option of the lineage-specific *LAVA* retrotransposon in the gibbon genome

**DOI:** 10.1101/765230

**Authors:** Mariam Okhovat, Kimberly A. Nevonen, Brett A. Davis, Pryce Michener, Samantha Ward, Mark Milhaven, Lana Harshman, Ajuni Sohota, Jason D. Fernandes, Sofie R. Salama, Rachel J. O’Neill, Nadav Ahituv, Krishna R. Veeramah, Lucia Carbone

## Abstract

Co-option of transposable elements (TEs) to become part of existing or new enhancers is an important mechanism for evolution of gene regulation. However, contributions of lineage-specific TE insertions to recent regulatory adaptations remain poorly understood. Gibbons present a suitable model to study these contributions as they have evolved a lineage-specific TE called *LAVA,* which is still active in the gibbon genome. The LAVA retrotransposon is thought to have played a role in the emergence of the unusually rearranged structure of the gibbon genome by disrupting transcription of cell cycle genes. In this study, we investigated whether LAVA may have also contributed to the evolution of gene regulation by adopting enhancer function. We characterized fixed and polymorphic LAVA insertions across multiple gibbons and found 96 LAVA elements overlapping enhancer chromatin states. Moreover, LAVA was enriched in multiple transcription factor binding motifs, was bound by an important transcription factor (PU.1), and was associated with higher levels of gene expression in *cis*. We found gibbon-specific signatures of purifying/positive selection at 27 LAVA insertions. Two of these insertions were fixed in the gibbon lineage and overlapped with enhancer chromatin states, representing putative co-opted LAVA enhancers. These putative enhancers were located within genes encoding SETD2 and RAD9A, two proteins that facilitate accurate repair of DNA double-strand breaks and prevent chromosomal rearrangement mutations. Thus, LAVA’s co-option in these genes may have influenced regulation of processes that preserve genome integrity. Our findings highlight the importance of considering lineage-specific TEs in studying evolution of novel gene regulatory elements.

## Introduction

Transposable elements (TEs) comprise nearly half of mammalian genomes and provide a major source of genetic and epigenetic variation during evolution. While many studies have focused on the disruptive consequences of TE insertions, especially those that impact human health (*1*), growing evidence is revealing the widespread presence of advantageous TE insertions across lineages (*2*). Depending on their impact on the host, TEs face different evolutionary fates. Most TE insertions are neutral and therefore drift randomly in the population, while disruptive insertions are actively selected against and removed from the population. The occasional adaptive TE insertions however, are favored and preserved by selection and may ultimately become incorporated into the host genome, in a process known as “co-option” or “exaptation”. To date, several examples of co-opted TEs have been reported across vertebrates [Reviewed in (*3*)].

Many co-opted TEs are capable of modifying gene expression in a tissue- or time-specific manner by forming new *cis*-regulatory elements (i.e. enhancers) or by being incorporated into already existing enhancers (*4*). Since regulatory TEs often contain transcription factor (TF) binding sites, their transposition in the genome can reshape entire gene regulatory networks by introducing similar regulatory modules nearby multiple genes (*5*). Furthermore, since the distribution of regulatory TE insertions often varies across lineages, co-option of these elements likely represents a major mechanism for evolution of lineage-specific gene regulation patterns (*5–7*). Indeed, a recent comparative study in primates demonstrated that nearly all human-specific regulatory elements overlapped TE sequences, and that most TE families enriched at *cis*-regulatory regions were relatively young and lineage-specific (*7*). These and other findings highlight young regulatory TEs as an important source for evolution of gene regulation in primates (*7–9*). However, due to technical challenges associated with studying recent TE insertions (e.g. low mappability), their contributions to the evolution of gene-regulatory adaptations still remains poorly understood, especially in non-human primates.

Among primates, the endangered gibbons (Hylobatidae) present an attractive model for exploring functional contributions of a lineage-specific TE. Gibbons (or small apes) occupy an important node in the primate phylogeny between old world monkeys and great apes; they have an intriguing evolutionary history, and have evolved many unique traits [e.g. locomotion via brachiating and monogamy (*10*)]. Most notably, the gibbon lineage has experienced drastic genomic rearrangements since its divergence from the common Hominidae ancestor ~17 million years ago (mya) (*11*). These evolutionary rearrangements are not only evident through comparisons with great ape genomes, but also in the vastly different karyotypes of the four extant gibbon genera: *Nomascus* (2n=52), *Hylobates* (2n=44), *Hoolock* (2n=38) and *Siamang* (2n=50), which split almost instantaneously around 5 mya. Factors leading to these evolutionary genome reorganizations are not fully understood, but a gibbon-specific retrotransposon called *LAVA* (Fig. 1A), may have played a role (*12*).

**Figure 1.**
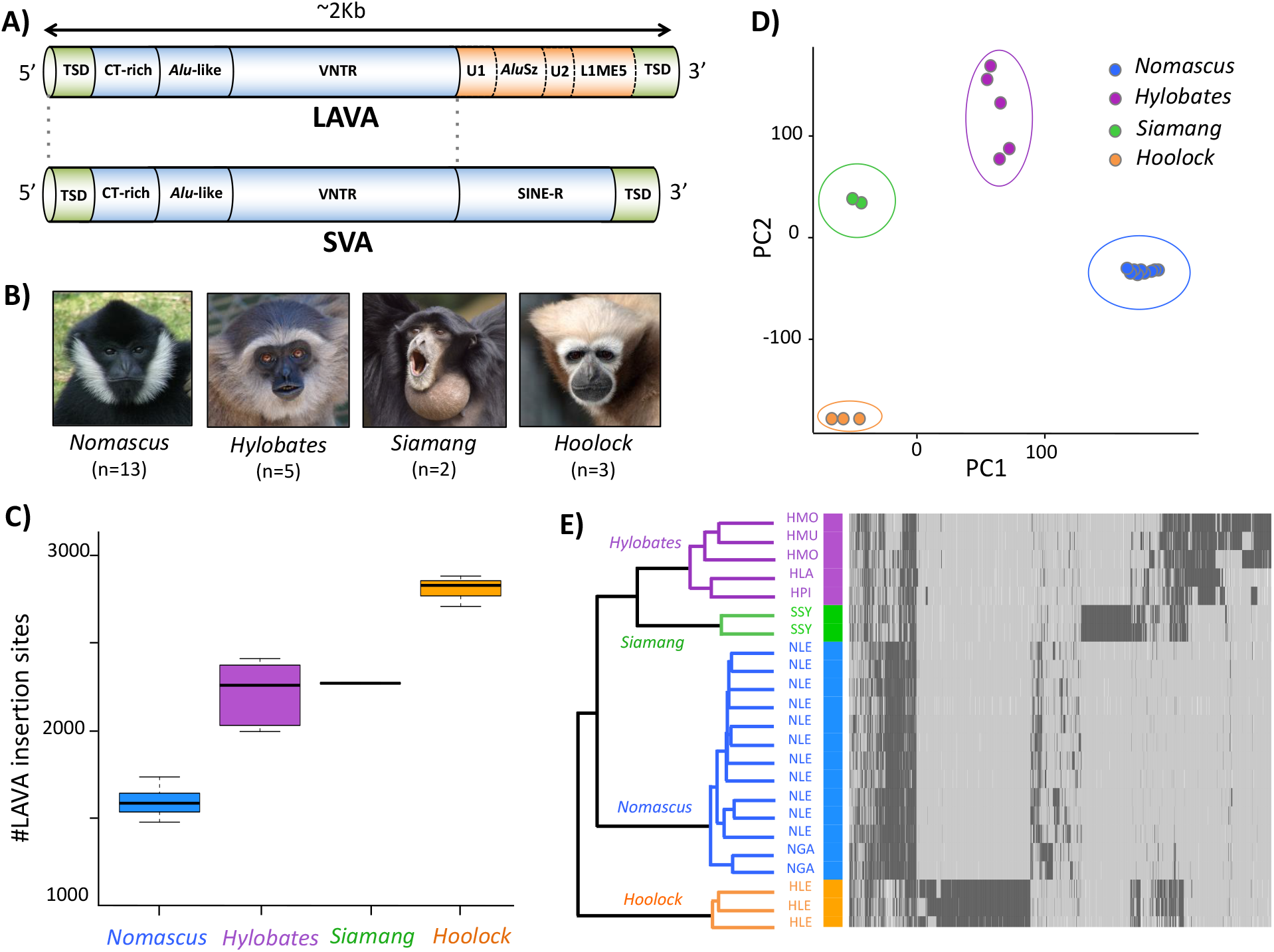
The LAVA element displays genus-specific expansion patterns. **A)** The full-length composite LAVA element shares structural components with SVA (TSD= target site duplication, VNTR= variable number tandem repeat, U= unique non-repetitive sequence). **B)** Representative species of the 4 extant gibbon genera shown along with our WGS sample sizes below each photo. **C)** The number of LAVA insertion loci per individual varied greatly across genera. **D)** Logarithmic PCA of LAVA genotypes groups individuals based on genus. **E)** Unsupervised hierarchical clustering of individuals, using LAVA genotype at 4,585 unfixed LAVA insertion loci, groups gibbons based on genus. In the heatmap, dark gray= presence of LAVA, light gray= absence of LAVA, and white= missing data. Species abbreviations are as follows: *Nomascus* [NLE= *N. leucogenys* and NGA= *N. gabriellae*), *Hylobates* (HLA= *H. lar*, HMO= *H. moloch*, HMU=*H. muelleri*, HPI= *H. pileatus*), *Siamang* (SSY= *S. symphalangus*), and *Hoolock* (HLE= *H. hoolock*)].

LAVA (LINE-*Alu*Sz-VNTR-*Alu*_LIKE_) is a non-autonomous retrotransposon that relies on the L1 protein machinery for its retrotransposition via target primed reverse transcription (*13, 14*). The full-length, ~2 Kb long composite LAVA element is found only in gibbons, but its structure is composed of portions of TEs commonly found across primate genomes, namely the 5’ portion of SVA (SINE-VNTR-*Alu*_LIKE_), as well as pieces of *Alu*Sz and L1ME5 elements (Fig. 1A). Despite sharing much of their overall structure, SVA and LAVA elements have had drastically different propagation success in the gibbon lineage. The SVA element, which has thrived in all great apes, is only present in ~30 copies in the gibbon genome (*15, 16*), whereas LAVA is estimated to be 20-40 times more abundant across gibbon genera (*12, 13*).

In the original analysis of the reference gibbon genome (Nleu3.0), which was derived from a northern white-cheeked gibbon (*Nomascus leucogenys*), nearly half of the >1,000 LAVA insertions found were located within or near genes, especially genes involved in regulation of cell cycle and chromosome segregation (*12*). Since some intronic LAVA insertions are capable of terminating gene transcription prematurely, disruption of cell cycle genes by LAVA was considered a contributing cause for the abundant evolutionary genomic rearrangements in the gibbon lineage (*12*). However, LAVA’s successful and ongoing propagation in the gibbon genome (*14*), in spite of its disruptive effects, may have allowed a subset of insertions to adopt adaptive functions. In fact, TEs that are prevalent near genes have a stronger propensity for adopting enhancer function and modulating expression of adjacent genes (*17*). Thus, adaptive contributions from LAVA may have involved regulation of nearby genes and might have ultimately resulted in its co-option as a *cis*-regulatory element in the gibbon genome.

In this study, we characterized LAVA insertions across multiple gibbon genomes and used genomic, epigenetic, and evolutionary analyses to investigate evidence of LAVA’s functionality and co-option in the gibbon genome.

## Results

### Genome-wide identification of LAVA insertion/deletions (indels) across gibbons reveals genus-specific expansion patterns

Since the LAVA element is still able to retrotranspose in the gibbon genome (*14*), its distribution is expected to vary across unrelated gibbons. To this end, we generated whole genome sequencing (WGS) datasets from 23 unrelated gibbons across the four extant genera (*Nomascus, Hylobates, Siamang and Hoolock*; Fig. 1B, Table S1). We selected the Mobile Element Locator Tool [MELT (*18*)] to identify TE indels from short-read data as this software was found to effectively identify SVA elements from human WGS datasets (*18, 19*). MELT uses discrepancies in WGS data aligned to a reference genome to identify non-reference TE insertions, as well as deletion of TEs present in the reference, and then combines information across datasets to characterize and genotype (presence/absence) TE indels in a population.

Through *in silico* simulation analyses, we first validated MELT’s capability to identify LAVA indels and showed that ≥10X WGS coverage is required for identifying ≥75% of LAVA indel sites within 10 bp of their true position with high sensitivity and specificity (Supplemental Text, Figs. S1). We then used MELT to identify LAVA indels across our 23 WGS datasets, which all had >10X coverage (Table S1). Since all WGS datasets were aligned to the same gibbon reference genome [Nleu3.0, generated from the *Nomascus leucogenys* (NLE) species (*12*)] we were able to use MELT to identify and genotype orthologous LAVA insertion loci across genera. We identified an initial list of 20,734 non-reference LAVA insertion sites that we combined with 1,118 LAVA insertions previously identified in the Nleu3.0 assembly (*12*). To minimize false positives, we filtered the total 21,852 LAVA insertions based on strict quality, length and frequency criteria, which reduced the total number of LAVA insertion sites to 5,490 high-confidence hits (Table S2). We found a significant effect of genus on the number of LAVA insertions identified (ANOVA, p<0.0001; Fig. 1C), as well as a significant effect of WGS coverage within each genus (p<0.0001). The smallest numbers of LAVA insertions were found in the *Nomascus* genus and the highest numbers were found in the *Hoolock,* the same genus in which LAVA was first discovered and found to form long centromeric expansions (*13, 20*).

Since LAVA is rarely deleted post-insertion (*12*), genotype differences at LAVA insertion loci are mostly due to differential insertions. Of all 5,490 LAVA insertion loci found, nearly half (2,888 or 52.6%) were found exclusively in some or all individuals from a single genus (genus-specific), indicating LAVA’s insertion at these sites likely occurred following the split of the four gibbon genera ~5 mya (*12*). In contrast, a quarter of insertion loci (1,317 or 24.0%) were discovered in at least one individual in each genus (i.e. shared), suggesting that LAVA insertion at these sites predated the genera split. Over half (905 or 68.7%) of these shared insertion loci were homozygous for the presence of LAVA in all 23 gibbons and were likely fixed in the gibbon lineage. Logarithmic principal component analysis (PCA) of genotypes at 4,585 unfixed LAVA insertions grouped individuals of the same genus together (Fig. 1D), and unsupervised clustering analysis organized gibbons in a pattern resembling a potential LAVA-based gibbon phylogeny (Fig. 1E). While this dendrogram recapitulates some of the published phylogenies obtained from gibbon mitochondrial DNA (*21, 22*), it does not match more recent phylogenies obtained by us and others based on nuclear genomes (*23, 24*). Therefore, it should be interpreted merely as a validation of our LAVA (presence/absence) genotyping pipeline, rather than a true gibbon phylogeny, which still remains elusive.

### LAVA insertions vary in population frequency and are unevenly distributed in the genome

To reduce errors from cross-species alignment of WGS datasets, we focused all downstream analysis on LAVA identified in *Nomascus leucogenys* (NLE) gibbons, the same species used to build the gibbon reference genome (*12*). Of the total 2,266 LAVA insertion loci found among NLE individuals, 48% (1,095) had homozygous LAVA insertions in all 11 NLE individuals. These LAVA insertions have reached high frequency or fixation due to selection or drift, and were thus called *fixed-LAVA.* The remaining 1,171 LAVA insertions sites were present in lower population frequencies (<95%), and displayed presence/absence polymorphism, hence they were called polymorphic or *poly-LAVA* (Fig. 2A, Table S2).

**Figure 2.**
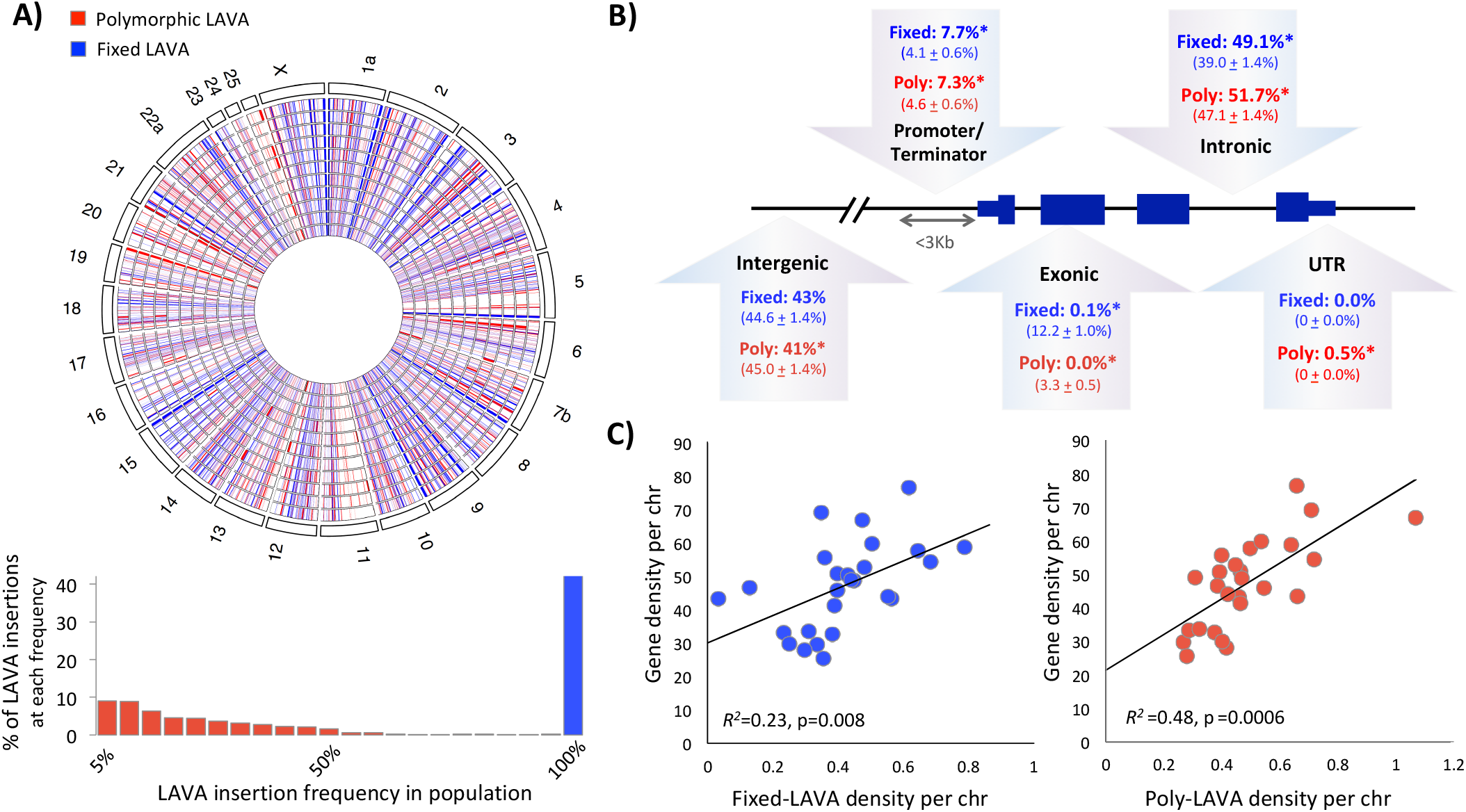
Fixed- and poly-LAVA insertions have similar and uneven genomic distribution. **A)** LAVA insertions present at 100% frequency in the population were classified as fixed-LAVA (blue), and the rest were classified as poly-LAVA (red). Circos plot (*top*) shows the non-homogenous distribution of fixed- (blue) and poly-LAVA (red) insertions across the gibbon chromosomes. Chromosome numbers are annotated on the outer circle, and 11 inner circles represent the genomes of the 11 NLE gibbons included in this study. **B)** Proportion of LAVA observed in different positions with respect to genes is shown in bold and mean± stdev expected percentages are reported below. (*) indicates significant (permutation p<0.01) deviation from expectation. **C)** Correlations between gene- and LAVA-density per chromosome are shown for fixed- (blue) and poly-LAVA (red).

In general, LAVA insertion loci were unevenly distributed across chromosomes (χ^2^_fixed-LAVA_ ^(25, N=1095)^ and χ^2^_poly-LAVA_ ^(25, N=1171)^, p<0.001; Figs. 2A, S2A) and formed broad clusters within chromosomes (one-tailed permutation p<0.001, Supplemental Text), likely due to genomic context such as repeat and gene density. Consistent with observations for other retrotransposons (*25*), LAVA insertions were enriched near repeats (permutation q<0.05; Fig. S2B). Furthermore, both fixed- and poly-LAVA insertion loci were significantly closer to genes than expected by random chance, and were over-represented in non-coding regions of genes (i.e. introns, promoters, or terminators; permutation p <0.001; Fig. 2B). In line with these observations, we also found significant correlation between LAVA- and gene-density across chromosomes (R^2^ _fixed-LAVA_ =0.23, p=0.008 and R^2^ _poly-LAVA_=0.48, p =0.0006; Fig. 2C). Hence, both fixed- and poly-LAVA insertion sites have uneven distribution in the genome and are over-represented near genes, likely as a result of LAVA’s preferential insertion into open chromatin (*26, 27*), effects of selection post-insertion, or a combination of both.

### Several LAVA elements show chromatin signatures of enhancer activity

To investigate if LAVA elements have evolved regulatory function in the gibbon genome, we sought to identify LAVA elements displaying epigenetic hallmarks of enhancer activity. We performed chromatin immunoprecipitation sequencing (ChIP-seq) against three activating (H3K4me1, H3K27ac, and H3K4me3) and two repressing histone marks (H3K27me3 and H3K9me3), using EBV-transformed lymphoblastoid cell lines (LCL) established previously (*12, 28*) and in this study, from three unrelated NLE individuals (Table S4, Supplemental Text). We annotated the epigenetic landscape of the reference gibbon genome using nine chromatin states, each of which represented a different combination of histone marks (Fig. 3A). Since chromatin state assignment is restricted to sequences present in the reference genome, we were only able to investigate and report chromatin states for the subset of LAVA insertions present in the Nleu3.0 assembly (all 1,095 fixed-LAVA, and 23 poly-LAVA). We calculated “fold-enrichment” of chromatin states overlapping these elements by accounting for length and prevalence of LAVA and chromatin states in the whole genome. Of states overlapping fixed- and poly-LAVA elements, constitutive “Heterochromatin” and “Polycomb-Repressed” states had the highest fold-enrichment, respectively (Fig. 3A). However, the chromatin state covering the most length (77% of fixed- and 75% of poly-LAVA; Fig. S4A) and number of LAVA elements (86% of fixed- and 87% of poly-LAVA; Fig. 3B) was Low Signal/Mappability, reflecting the difficulty in aligning short reads to repetitive sequences. Of note, we identified several LAVA elements that overlapped enhancer chromatin states; specifically, 13 fixed-LAVA overlapped bivalent enhancer chromatin, 72 fixed- and one poly-LAVA colocalized with poised enhancer chromatin, and 23 fixed-LAVA overlapped with active enhancer states (Fig 3C). In total, 95 (8.7%) fixed LAVA insertions and one (4.3%) poly-LAVA element colocalized with at least one enhancer chromatin state. These overlaps ranged from 10-1,474 bp in length (median=182 bp), consistent with the typical length of enhancers [i.e. 10-1000 bp (*29*)], and comprised ~3.5% of the total length of all LAVA sequences in the reference genome (Fig. S4A).

**Figure 3.**
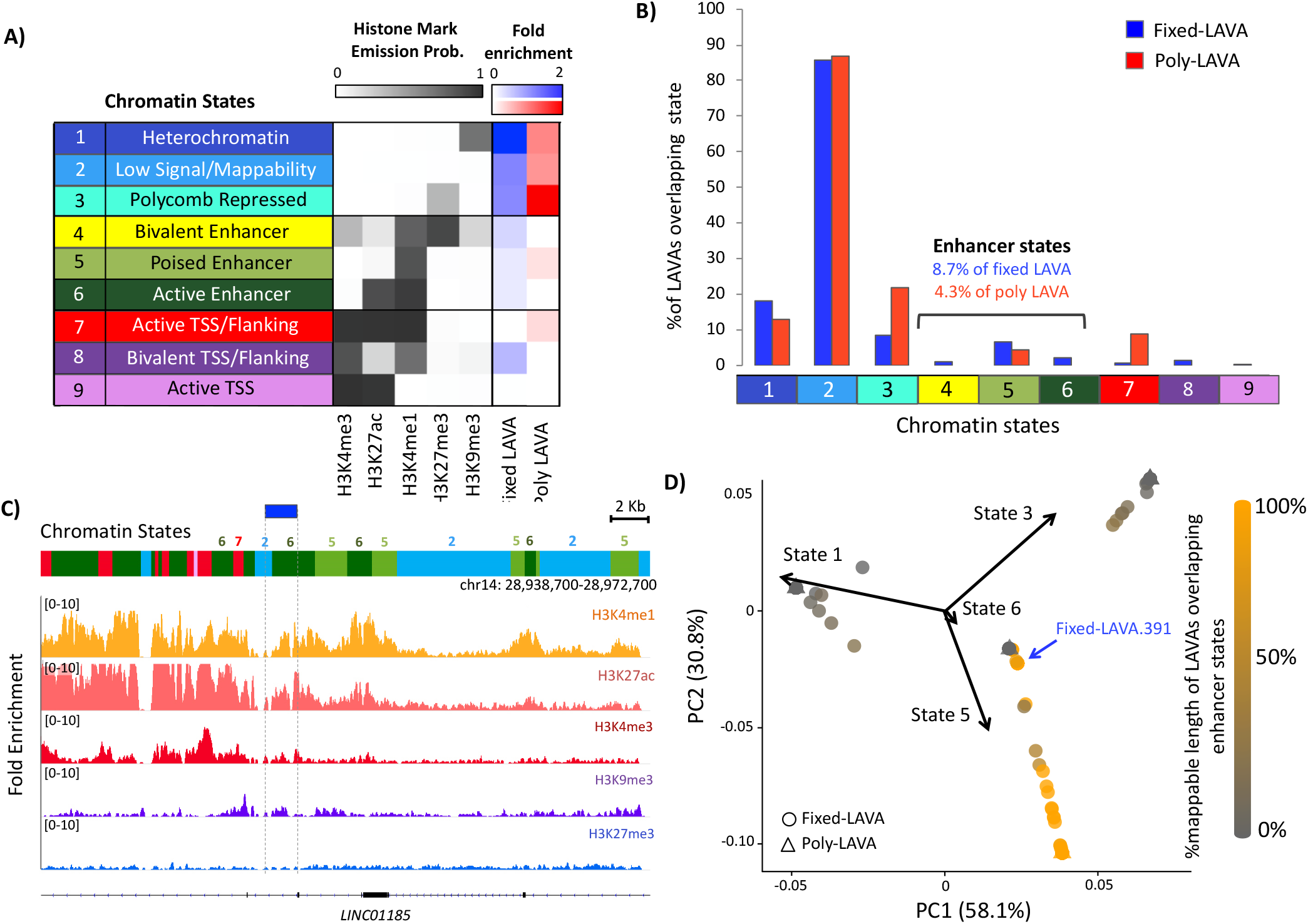
Several LAVA elements display epigenetic signatures of enhancers. **A)** Chromatin states were characterized based on histone ChIP-seq signal. Fold enrichment of states overlapping reference-present fixed- (blue) and poly-LAVA (red), relative to random expectation, is shown. **B)** Percent LAVA insertions with ≥10 bp overlap with each chromatin state are shown. X-axis colors and numbers correspond to chromatin states from panel A. **C)** Example of a fixed-LAVA that overlaps Active Enhancer chromatin state. ChIP-seq fold-enrichment tracks (relative to INPUT) are shown below the chromatin states. **D)** PCA biplot of 393 LAVA elements based on chromatin state compositions. LAVA elements are shaded based on % of their length overlapping with enhancer states (states 4, 5 or 6), after excluding overlap with Low Signal/Mappability chromatin state.

PCA analysis of LAVA chromatin state composition (excluding overlaps with Low Signal/Mappability chromatin state and the 725 LAVA element entirely covered by this state) grouped LAVA into three epigenetically distinct groups; one group consisted mostly of LAVA overlapping enhancer states (Fig 3D), while the other two groups consistent of putatively silenced LAVA elements colocalizing with either Constitutive or Polycomb Repressed chromatin (Fig S4B,C).

### LAVA elements provide transcription factor binding motifs and some elements are bound by PU.1

Regulatory TEs contribute roughly 20% of all transcription factor (TF) binding sites in mammalian genomes (*30*). Using two different pipelines, we identified a conservative list of six TFs whose recognition motifs were significantly overrepresented (q<0.05) in LAVA sequences and were predicted to bind LAVA with high affinity: PU.1 (encoded by *SPI1*), STAT3, SRF, SOX10, SOX17, and ZNF143 (Fig S5A). Since motif enrichment analysis on highly similar and repetitive sequences may lead to false positives, we sought to experimentally validate binding of one candidate TF to LAVA. We focused on PU.1, an important TF in the development of B-lymphoid cells (*31*), whose recognition motif was highly enriched in LAVA, especially in the VNTR region (Figs. 4A, B and S5B). To validate PU.1 binding to LAVA, we performed ChIP-seq against PU.1 in two gibbon lymphoblastoid cell lines (Table S4). We first used the RepEnrich2 software (*32*), which takes advantage of both unique and multi-mapping sequencing reads to assess overall enrichment of repeat families in ChIP-seq samples relative to input (indicating binding), without the need for high mappability or peak-calling. Of the 16 repeat families significantly enriched in gibbon PU.1 ChIP-seq samples, the “SVA repeat family”, which is composed of SVA and LAVA insertions, was the most significantly enriched (log fold change= 0.4, q=1.52e-59; Fig. 4C). It should be noted that in gibbons, enrichment of “SVA repeat family” can be equated to enrichment of LAVA repeats, because (with only ~30 insertions genome-wide) SVA elements make up a negligible proportion of this family (*15, 16*).

**Figure 4.**
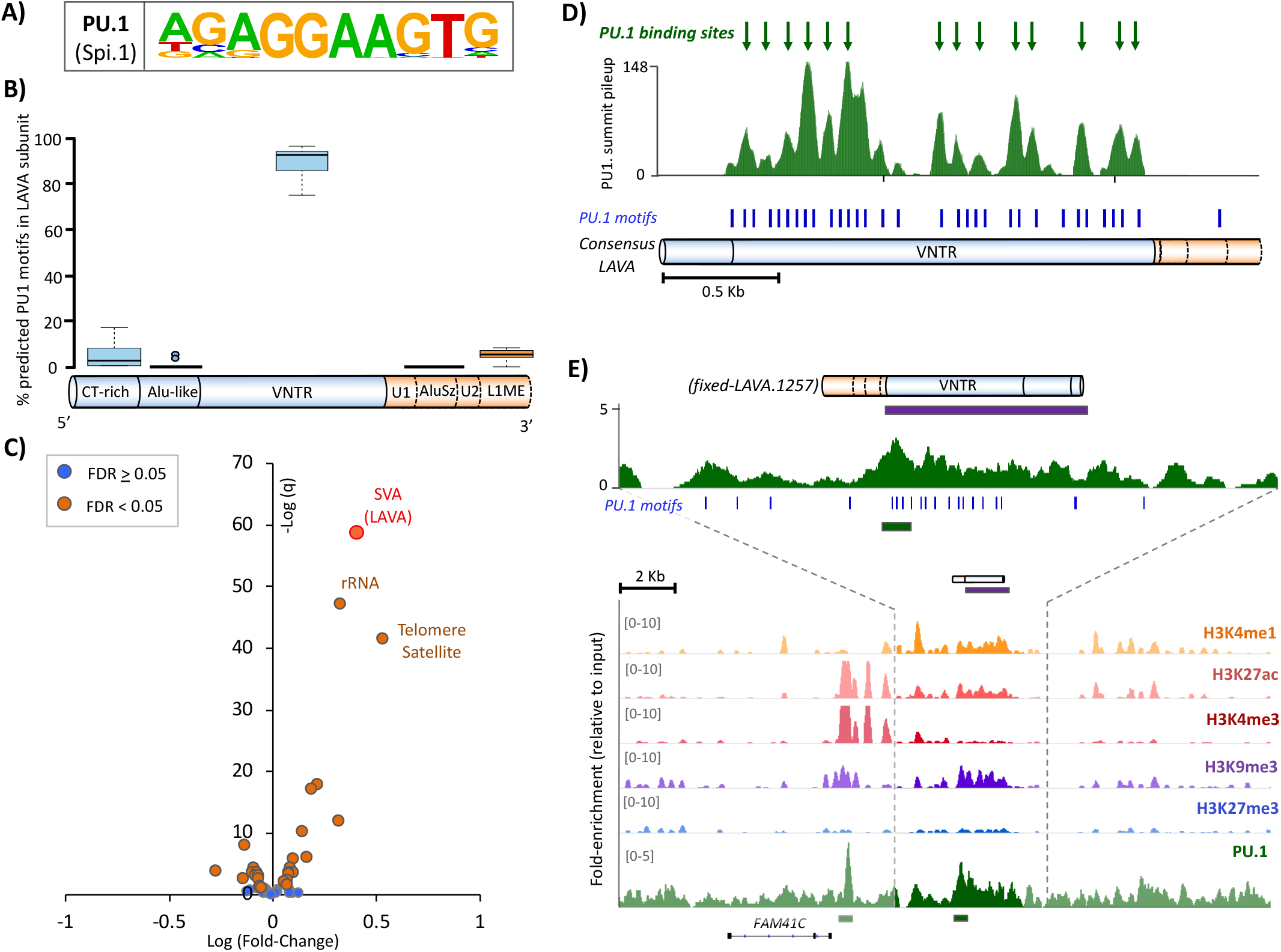
The PU.1 transcription factor binds LAVA at the VNTR subunit. **A)** PU.1 consensus binding motif. **B)** Proportions of LAVA’s predicted PU.1 binding motifs found in each subunit (based on 20 randomly selected full-length LAVA elements.) **C)** Volcano plot displays enrichment of repeat families in gibbon PU.1 ChIP-seq. **D)** PU.1 ChIP-seq summit pileups are shown along the full-length LAVA consensus. Putative PU.1 binding sites (i.e. meta-summits) are marked with green arrows. Blue ticks indicate predicted PU.1 binding motifs. **E)** A screenshot of the fixed-LAVA containing a significant PU.1 ChIP-seq peak (green bar) and overlapping bivalent enhancer chromatin state (purple bar). Predicted PU.1 binding motifs are marked with blue ticks.

To characterize sites of PU.1 binding within LAVA elements, we used multimapping and uniquely aligning reads to identify 22,264 peaks between our two replicates. Of these, the apex (i.e. summit) of 138 peaks overlapped LAVA elements present in the reference gibbon genome. Using an approach similar to Fernandes et al. (*33*), we marked the positions of these overlapping summits within a consensus full-length LAVA element to generate a pileup of summit positions (Fig. 4D). Apexes of this pileup (i.e. meta-summits), which represent putative PU.1 binding sites, were all located inside the VNTR subunit and in overall agreement with the distribution of *in silico* predicted PU.1 recognition motifs (Fig. 4D). Lastly, to localize specific LAVA insertions binding PU.1 in the gibbon genome, we performed peak-calling only using uniquely mapping reads and identified 13,920 unique peaks, which were collectively enriched for the consensus PU.1 recognition motif and co-occurred with active histone marks (Fig. S5C-D). In this approach, most overlaps between PU.1 peaks and LAVA are expected to be missed due to removal of multi-mapping reads, nonetheless we found significant PU.1 peaks inside the VNTR of two fixed-LAVA elements. Based on our previous chromatin state analysis, one of these elements (fixed-LAVA.1257) overlapped bivalent enhancer chromatin state (Fig. 4E) while the other (fixed-LAVA.1087) was entirely covered by Low Signal/Mappability chromatin state.

Since PU.1 appears to bind LAVA at the VNTR, a subunit shared between LAVA and SVA elements (Fig. 1A), we investigated whether the ability to bind PU.1 was specific to LAVA or shared with SVA. After repeating our analysis on public human ENCODE data, we did not detect any enrichment of the “SVA repeat family” in human PU.1 ChIP-seq datasets [q=0.8; Fig. S6A; (*34, 35*)]. We also did not find any PU.1 ChIP-seq summits mapping to the consensus human SVA sequence. Consistently, comparison of PU.1 recognition motifs between LAVA and SVA sequences revealed that: 1) the consensus SVA sequence contained fewer PU.1 recognition motifs compared to LAVA (5 vs. 32; Figs.S5B and S6B), 2) a lower proportion of SVA repeats in the human reference genome contained PU.1 motifs (56% of SVAs vs. 41% of random size-matched background sequences) compared to LAVA in the gibbon genome (100% of LAVAs vs. 56% in background), and 3) the average density of PU.1 motifs in SVA repeats (2.5 ± 4.2; mean±stdev motifs per 1 Kb) was lower compared to LAVA (8.6± 4.1; Mann-Whitney-Wilcoxon Test, p-value < 2.2e-16). Thus, despite the close relationship between LAVA and SVA, major differences exist in their ability to bind PU.1, and potentially other TFs, likely as a result of sequence differences that have evolved in the VNTR region since LAVA diverged from SVA in the gibbon lineage (*36*).

### Genes near LAVA have overall higher expression compared to the rest of the genome

To investigate LAVA’s potential effect on expression of nearby genes, we generated RNA-seq data from nine NLE gibbon LCLs generated previously (*12, 28*) and in this study. We considered genes with depth normalized read counts (count per million, CPM) higher than 0.5 in at least two of the nine gibbons to be “actively expressed”. Using this proxy, 72% of the nearest genes within 3 Kb of fixed-LAVA (448 out of 620) and 69% of the nearest genes within 3 Kb of poly-LAVA (478 out of 694) were considered actively expressed. These proportions were not significantly different from each other (two tailed Fisher’s exact test, p=0.58), but were both significantly higher than the 39% of genes (15,715 out of 40,504) that were actively expressed genome-wide (two-tailed chi-square test with Yates correction, p<0.0001). Moreover, among actively expressed genes, those located near fixed- and poly-LAVA had significantly higher median expression compared to the null distribution in the whole-genome (two-tailed permutation test, p<0.0001; Fig. 5A).

**Figure 5.**
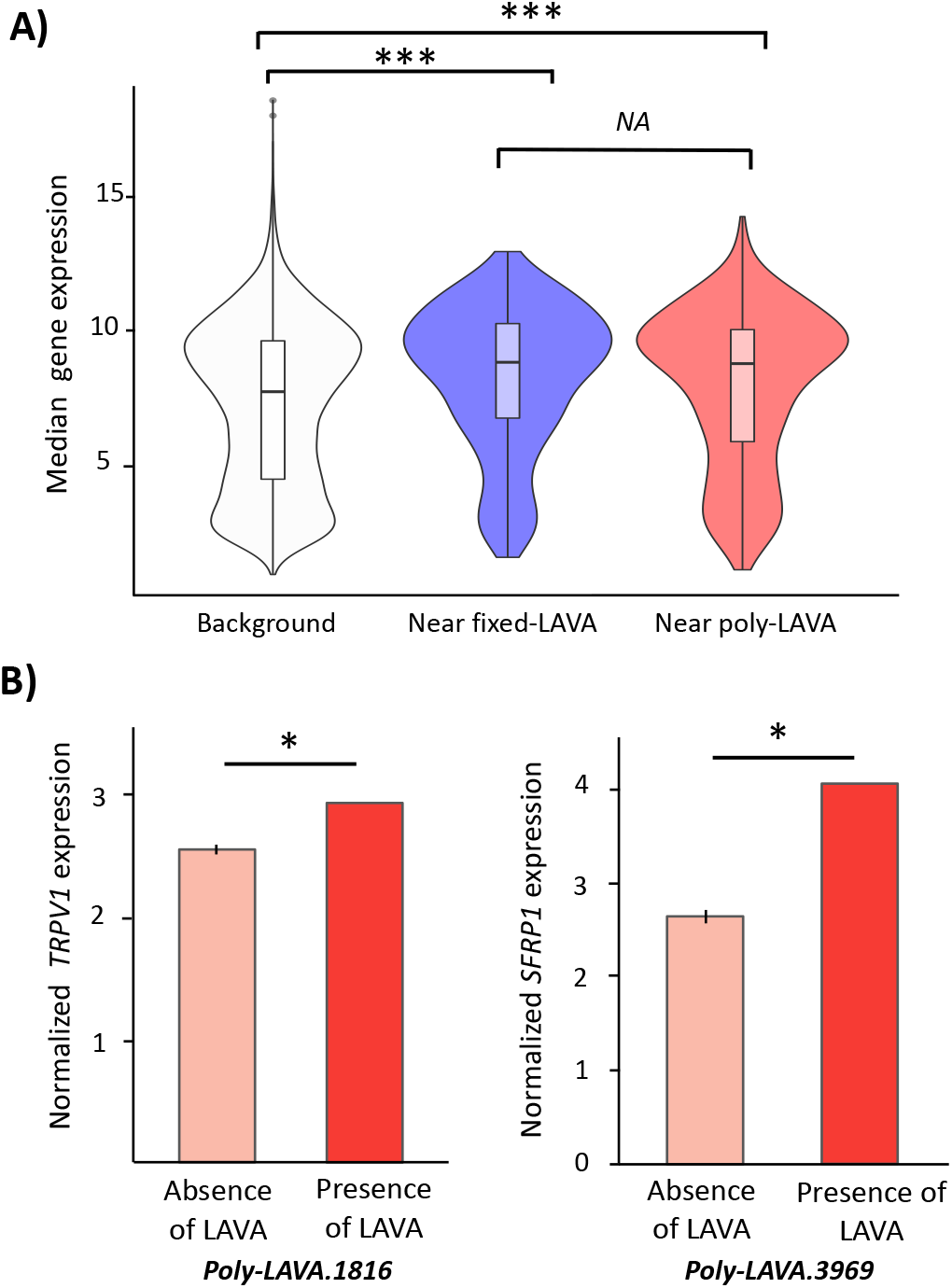
LAVA is associated with higher expression of genes in *cis*. **A)** Violin plots demonstrate distribution of median normalized gene expression for genes not located near LAVA insertions (white) vs. those ≤3 Kb of fixed- (blue) and poly-LAVA (red) (***= permutation p<0.0001). **B)** Normalized expression of *TRPV1* (*left*) and *SFRP1* (*right*) are shown against genotype at corresponding poly-LAVA. Bar heights represent mean normalized expression and error bars are standard deviation (*= q<0.05). All Y-axis are rlog transformed gene counts normalized for gene length, GC content and sequencing depth (i.e. log2 scale).

Next, we took advantage of the presence/absence of poly-LAVA insertions and examined correlation between LAVA genotype and expression level of genes within 1Mb of LAVA insertion loci. Despite being underpowered due to our small sample-size of nine, we found two poly-LAVA insertions associated with significant increase in expression of a nearby gene (q<0.05; Fig. 5B). One of these genes was *TRPV1* (transient receptor potential cation channel subfamily V member 1), whose expression was significantly higher in an individual with a LAVA insertion ~300 Kb downstream (p=8.38e-07, q=0.01; Fig. 5B, left). In humans, *TRPV1* is highly expressed in the central nervous system and has a human-specific SVA insertion hypothesized to have contributed to the evolution of human-specific behaviors (*37*). The other gene encodes the secreted frizzled related protein (SFRP1), a Wnt antagonist implicated in cell-cycle regulation and senescence (*38, 39*), which was more highly expressed when a LAVA insertion was present ~800 Kb upstream (p=3.8e-06, q=0.04; Fig. 5B, right). While our correlative analyses are consistent with these poly-LAVA elements functioning as *cis*-regulatory elements, neither insertion was present in the reference genome and their chromatin state could not be determined.

### Several LAVA insertions show signatures of gibbon-specific selective constraint

To investigate co-option of LAVA, we examined LAVA insertion loci for evidence of selection. We used Tajima’s D (*40*), a summary statistic that measures difference between two estimates of population genetic diversity (θ). Significantly negative Tajima’s D values indicate an excess of low frequency alleles compared to expectation, compatible with a recent selective sweep or background selection. We measured mean Tajima’s D in two 10 Kb windows directly flanking each side of LAVA insertion sites and found that 11/808 poly-LAVA and 19/734 fixed-LAVA elements included in our evolutionary analysis had Tajima’s Ds within the 5% most negative values in the genome (i.e. p<0.05 based on an empirical distribution of 10 Kb pairs genome-wide). These loci had Tajima’s values <-2.0, compared to a genome wide estimate of −0.95, suggestive of selection occurring on either side of, and presumably within, these LAVA elements. The most noteworthy was a ~300 Kb cluster of four fixed-LAVA insertions on chromosome 18, which overlapped a major dip in Tajima’s D and displayed some of the lowest Tajima’s D values in the whole genome (Fig. S7). We also fit a demographic model to the NLE allele frequency spectra at a set of putatively neutral loci using ∂a∂i (*41*) and generated coalescent simulations under neutrality, to estimate a simulation-based p-value for each LAVA element. With the exception of two of the fixed-LAVA loci, all fixed- and poly-LAVA with p<0.05 based on the empirical criteria described above, also had p<0.001 based on this simulation framework, and p<0.05 even under the most conservative simulation regime of no recombination, which artificially decreases null Tajima’s D values (*42*).

If the selection signals identified in gibbons were independent of LAVA, for example, if they merely reflected background purifying selection due to proximity to genes, we may expect orthologous sequences in a sister taxon to show similar selection signals. Therefore, we examined genetic diversity in humans within orthologous regions to the windows flanking LAVA elements. We successfully identified orthologous regions for 8 (of the 11) poly-, and 13 (of the 17) fixed-LAVA elements that displayed significant signatures of selection in gibbons (under both empirical and conservative simulation frameworks). Of these orthologous regions, none had significant Tajima’s D values in human, with the exception of one poly-LAVA. In total, we found 10 poly- and 17 fixed-LAVA that showed significant signatures of positive/purifying selection in gibbon, but not in human (when orthologs were found), suggesting they, or their immediately surrounding regions, have acquired gibbon-specific functional properties.

### Enrichment and potential co-option of a subset of LAVA insertions near DNA repair genes

By investigating genes located near LAVA insertions, we can identify candidate biological processes influenced by LAVA. While no significant gene ontology (GO) term was enriched among genes near (≤3 Kb) poly-LAVA, those near fixed-LAVA insertions displayed significant enrichment (q<0.1) for 15 biological functions, all related to DNA repair (e.g. double- and single-strand break repair), and five cellular components important in cell cycle (e.g. spindle-pole centrosome; Fig. 6A, Table S6). Enrichment of these GO terms was validated using permutation analyses that accounted for gene length and LAVA’s preferential insertion near genes (two-tailed permutation p<0.001; Supplemental Text). Overall, these results recapitulated the previously described association of LAVA with cell cycle and chromosome segregation genes (*12*). However, by characterizing new LAVA insertions across multiple individuals and classifying them based on frequency in the population, we were able to unravel a novel association between fixed-LAVA and DNA repair pathways.

**Figure 6.**
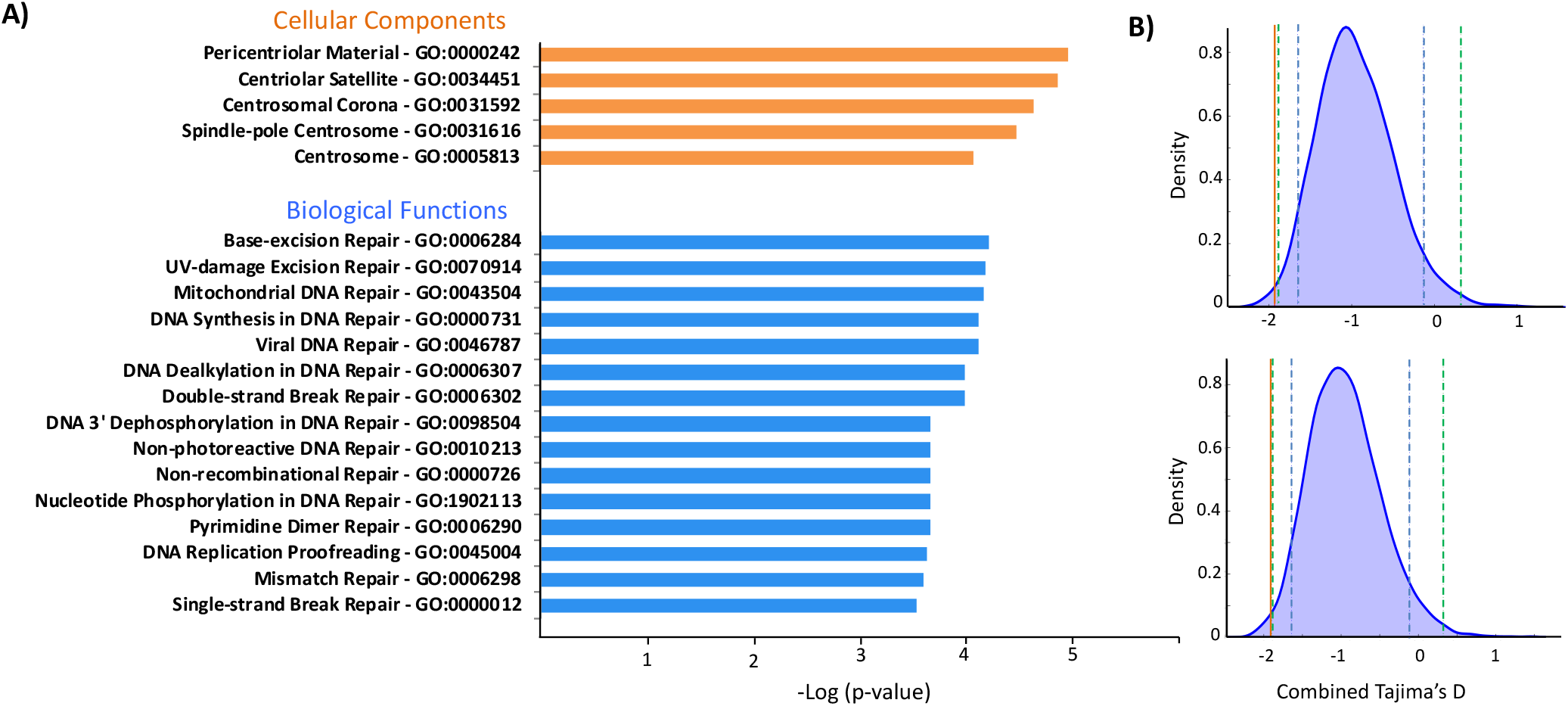
DNA repair genes are overrepresented near fixed-LAVA and putatively co-opted elements. **A)** Significant gene ontology terms among genes ≥3 Kb of fixed-LAVA are shown. The -log p-values are reported by Enrichr. **B)** The (up/downstream) combined Tajima’s D values of two putatively co-opted fixed-LAVA enhancers are shown (orange lines) against the corresponding genome-wide null distributions (blue bell curves). Dashed blue and green lines mark the 95^th^ and 99^th^ percentiles of the null distribution (see Supplemental Text for details).

We then searched for co-opted regulatory LAVA insertions, which are expected to be: 1) fixed in the gibbon genome, 2) show signatures of selection, and 3) overlap enhancer chromatin state. Based on these criteria, we found two putatively co-opted regulatory LAVA elements that were fixed across gibbon genera, had some of the smallest Tajima’s D values in the NLE genome (Fig. 6B) and overlapped poised enhancer state in gibbon LCL. These two LAVA insertions were located within introns of DNA repair genes encoding SETD2 (a histone methyltransferase) and RAD9A (a cell cycle checkpoint protein), which are specifically involved in facilitating accurate repair of DNA double-strand breaks (*43–45*). Considering the enrichment of several TF binding motifs in LAVA sequences, (for example, that of STAT3 (Fig. S5A), a major regulator of the DNA repair response), co-opted LAVA in these genes might be under selection to facilitate binding of TFs and alter regulation of DNA repair response globally or in a tissue/time-specific context.

Four other LAVA elements were also located in genes implicated in the timely detection and precise repair of DNA lesions [*TET3* (*46, 47*), *KMT2C* (*48*), *NSD2* (*49*) and *SMARCB1* (*50*)]. However, these LAVA elements did not overlap enhancer chromatin states in gibbon LCL, and may perhaps function in other developmental/tissue contexts or via other mechanisms.

## Discussion

In this study, we characterized insertions of the gibbon-specific LAVA retrotransposon across gibbon genera using the largest set of whole-genome sequencing data generated from these endangered species, to date. We found epigenetic and evolutionary evidence for the functionality of several LAVA insertions and showcased two putative co-opted LAVA enhancers within genes implicated in the accurate repair of DNA double-strand breaks.

We focused our investigation of LAVA’s functionality and co-option on the northern white-cheeked gibbon (*Nomascus leucogenys*, NLE), as this is currently the only gibbon species with a published reference genome (*12*). Among the subset of fixed- and poly-LAVA insertions that were present in the NLE reference genome, we found 96 unique elements overlapping active, poised, or bivalent enhancer chromatin states in gibbon lymphoblastiod cell lines (LCL). Consistent with known characteristics of enhancers (*51*), LAVA sequences were enriched in TF binding motifs including that of PU.1, an important regulator of gene expression in lymphocytes. After validating binding of PU.1 to LAVA in gibbon LCL, we showed that binding was mainly confined to the VNTR subunit, which harbors dense clusters of PU.1 binding motifs. Although LAVA’s VNTR originates from SVA (Fig.1A), PU.1 did not appear to bind SVA elements in the human genome. This finding was consistent with previous reports that LAVA’s VNTR has evolved distinct genetic features following its divergence from the SVA element (*36*). Furthermore, it indicated that LAVA has acquired unique epigenetic and functional properties, which might also be responsible for its successful propagation in the gibbon lineage (*14, 16*). Overall, we demonstrated that insertion of LAVA can introduce clusters of new (or additional) TF binding sites, which may alter regulation of nearby genes (*52, 53*). Notably, genes near LAVA had overall higher expression compared to the rest of the genome. Also, despite low statistical power, we found two positive correlations between presence of poly-LAVA loci and expression of a gene in *cis*. It should be noted however, that these observations were correlational and may, at least partially, reflect LAVA’s preferential insertion in actively transcribed chromatin.

Since biochemical activity of TEs could reflect selfish strategies for their survival and propagation rather than adaptive function (*27*), we also investigated signatures of selection at LAVA insertion sites. We identified significant signals of positive or purifying selection at 27 LAVA elements in gibbon, but not in any of the available orthologous regions in human, suggesting the selection signal is specific to LAVA elements. Two of these evolutionarily significant fixed-LAVA insertions also overlapped with poised enhancer chromatin state in gibbon LCL, representing two putative co-opted *cis*-regulator LAVA elements. Considering these elements overlap with poised enhancer states, and knowing that TE-derived enhancers often have tissue-specific activity (*4*), we predict that these putative co-opted LAVA enhancers are active in tissue/developmental contexts other than LCL. We also expect the true number of functional and co-opted LAVA insertions across gibbon tissues and genera to be larger than those identified here, considering all the challenges that limited our analysis, for example, unavailability of high-quality genome assemblies from all genera, exclusion of many LAVA elements from our epigenetic or evolutionary analyses, low mappability of short-read sequences to LAVA (*54*) and limited availability of tissues from the endangered gibbons.

While disruptive LAVA insertions are thought to have contributed to the emergence of genomic rearrangements in gibbons (*12*), evolutionary contributions of functional LAVA elements are not yet clear. Nonetheless, genes located near co-opted LAVA may provide some insight. The two putative co-opted *cis*-regulatory LAVA elements identified in this study were located in the introns of *SETD2* and *RAD9A* genes, both of which are crucial genes in maintaining genome integrity. SETD2 is a H3K36 histone methyltransferase that modifies chromatin at the site of DNA double-strand break to ensure its faithful repair via homologous recombination, rather than error-prone non-homologous end-joining (*43, 55*). RAD9A is a cell cycle checkpoint control protein that facilitates homologous recombination repair and prevents cell-cycle progression before DNA double-strand breaks are repaired (*44, 45*). Regardless of how or in which context the co-opted LAVA insertions may alter regulation of these genes, any adaptive regulatory changes resulting in improvement of DNA repair and genome integrity, would be favored/preserved by natural selection. As computational tools for studying TEs improve (*56*), and gibbon pluripotent stem cells (iPSC) provide access to currently unavailable tissues, we will be able to further investigate functional roles of co-opted LAVA across tissues, especially in the context of DNA double-strand repair. Insights from this and future studies will advance our understanding of how young TEs can contribute to lineage-specific evolution of gene regulatory novelty.

## Materials and Methods

See Supplemental Text for further details on most of the section described below.

### Genome-wide identification, genotyping and characterization of LAVA insertions

Genomic DNA extracted from blood of 23 unrelated gibbons across the four extant genera (*Nomascus*= 13, *Hylobates*= 5, *Hoolock*= 3, *Siamang*= 2) was used to construct whole genome sequencing (WGS) libraries as described before (*12*) and sequenced on Illumina HiSeq2000 and HiSeq2500 platforms. All gibbon WGS data were aligned using BWA (*57*) to the gibbon reference genome (Nleu3.0). MELT v2.1.3 (*18*) was used to predict non-reference LAVA insertions and deletions (indels) from WGS alignments, similar to our simulation analysis (see Supplemental Text). LAVA elements were annotated based on their insertion position relative to closest gene: intergenic, exonic, intronic, promoter (<3 Kb upstream of gene) or terminator (<3 Kb downstream of gene). LAVA predictions were filtered to remove: 1) low-quality inserts (as determined by MELT). 2) insertions present in only one copy in the population of 23 diploid genomes (i.e. heterozygous presence in only one genome), 3) insertions shorter than 290 bp (the minimum length required to discriminate a composite LAVA from its non-SVA subunits), and 4) inserts found on unplaced Nleu3.0 contigs. We also generated binary LAVA genotype profiles for all gibbon individuals (heterozygous or homozygous LAVA insertion=1, homozygous absence of LAVA= 0) and performed hierarchical clustering using the *hclust* function with the ward D2 method in R 3.6.1 and visualized the results with *heatmap.2* function in the *ggplot2* package. Logistic principal component analysis was carried out using the *logisticPCA* package in R (k=2 and m=4).

To reduce erroneous LAVA predictions due to cross-species WGS alignment, we only used LAVA insertions identified in 11 NLE individuals for downstream analysis. LAVA insertions found in two copies in all NLE individuals were called fixed-LAVA, while the rest were called polymorphic LAVA (poly-LAVA). Due to sequence ambiguity and absence of many poly-LAVA insertions from the reference genome, we did not include sequence polymorphism in our characterizations. We used permutation tests to characterize LAVA’s distribution across and within chromosomes, and relative to repeats and genes (see Supplemental Text for details).

### Histone chromatin immunoprecipitation sequencing (ChIP-seq) and chromatin state characterization

ChIP-seq was performed on gibbon EBV transformed lymphoblastoid cell lines (LCLs) from three unrelated NLE individuals, as previously described (*12*) and outlined in the Supplemental Text. Raw reads were QC’d with FastQC (*58*). All reads were aligned to Nleu3.0 using BWA (*57*) with default single-end settings, and low-quality/multi-mapping read alignments (MAPQ<30) were removed using samtools. ChIP-seq replicates displayed high correlation [Pearson correlation coefficient 75-96%; Fig. S3; (*59*)], and were therefore combined (Fig. S3). ChromHMM (*60*) was used to identify and characterize 9 chromatin states based on the histone ChIP-seq alignments. “Fold enrichment” of each chromatin state at reference-present LAVA was calculated as (C/A)/(B/D), where: “A” is the genome-wide number of bases in the state, “B” is the number of genome-wide bases in LAVA, “C” is the number of bases in the state and LAVA, and “D” is the number of bases in the genome. Lastly, we used BEDtools (*61*) to characterize overlap of chromatin states with reference-present LAVA elements, requiring each overlap to be ≥10 bp long. For each LAVA, we calculated percentage of its length overlapping different chromatin states after excluding regions overlapping Low Signal/Mappability. We also removed 725 LAVA elements that had 100% overlap with Low Signal/Mappability state. Chromatin compositions of the remaining 393 elements were used to perform PCA analysis in R.

### Transcription factor motif enrichment and PU.1 binding to LAVA

Sequences of reference-present LAVA insertions on the assembled chromosomes of Nleu3.0 were used to perform motif enrichment analyses with the Homer suite (*62*), and the Transcription factor Affinity Prediction [TRAP; (*63*)] web tool. To meet length restrictions, we removed 24 LAVA sequences longer than 3 Kb in TRAP analysis. P-values were corrected using the Benjamini-Hochberg method (*64*) and only significant motif enrichments (q<0.05) that agreed between the two methods were considered (Table S3). LAVA and SVA_A consensus sequences were obtained as described in the Supplemental Text, and their PU.1 motifs were predicted using the Homer suite (*62*). To assess distribution of PU.1 motifs across subunits of LAVA, we randomly selected 20 full-length LAVA elements, predicted PU.1 motifs using the Homer suite, and determined percentage of motifs found in each subunit. PU.1 ChIP-seq was performed on two gibbon LCLs, analyzed and compared to public human LCL PU.1 ChIP-seq data from ENCODE [GEO accessions GSM803531 and GSM803398; (*34, 35*)], as described in the Supplemental Text.

### Characterization of gene expression patterns near LAVA

RNA-seq gene count data was collected from two previously- (*12, 28*)and seven newly-established NLE LCLs, and normalized as described in Supplemental Text. Linear regression between LAVA (presence/absence) genotypes at each poly-LAVA locus and expression of genes within 1Mb, was performed using Matrix-eQTL with default settings (*65*). We used the GraphPad tool (https://www.graphpad.com/quickcalcs/contingency1.cfm) to perform a two-tailed chi-squared test with Yates correction and compare the proportion of “active genes” nearby fixed- and poly-LAVA to the rest of the genome. We used custom R scripts described in Supplemental Text to compare median of median expressions for genes located near LAVA to the rest of the genome.

### Assessing selection around LAVA insertion sites

A detailed description of evolutionary analyses is available in the Supplemental Text. Briefly, we used ANGSD (*66*) to estimate folded allele frequency spectra and Tajima’s D in 10 Kb windows. After filtering LAVA elements based on WGS coverage, we compared Tajima’s D in 10 Kb windows upstream and downstream of these elements to randomly sampled loci in the genome, and generated empirical p-values. We considered a LAVA element under selection only if p<0.05 for both upstream and downstream regions. In our simulation analyses, we generated simulated p-values by comparing observed Tajima’s D in 20 Kb windows centered at LAVA insertions to expectation under neutrality. To generate comparative human data, alignments from 20 Yoruba individuals from the 1000 Genomes Project (*67*) were used to calculate genome-wide Tajima’s D. We then used the multiz100way alignment to find orthologous human regions (Hg19) for the 10 Kb windows flanking LAVA elements and assessed significance of Tajima’s D in these regions using the empirical framework used for gibbons.

### Gene ontology (GO) analysis of genes nearby fixed- and poly-LAVA

We used Enrichr (*68*) with GO_Biological_Process_2017_7b, GO_Cellular_Component_2017_7b and GO_Molecular_Function_2017_7b libraries to test GO enrichment among the nearest genes (within ≤ 3 Kb) of fixed- and poly-LAVA. Significance of all GO terms that had p<0.05 and q<0.1 was validated using two permutation tests described in the Supplemental Text.

## Data availability

All WGS, ChIP- and RNA-seq data is available at Gene expression Omnibus (GEO) under accession no. GSE136968. Gibbon gene annotation file will be provided upon request.

## Acknowledgements

Authors would like to thank the zoos (San Antonio Zoo and Aquarium, Point Defiance Zoo and Aquarium, Oregon Zoo, Gladys Porter Zoo and Los Angeles Zoo) and the staff at the Gibbon Conservation Center (Santa Clarita, CA), especially the director Gabriella Skollar, who have provided us with opportunistic gibbon blood samples. The authors would also like to thank Drs. Jeff Wall and Michael Hammer for their invaluable contributions to WGS data collection, Dr. Eugene Gardner for aiding in optimizing the MELT pipeline, Dr. Jessica Minnier for guidance in RNA-seq analysis, Dr. R. Alan Harris, Patty Langasek and Christopher Klocke for their help with data analysis, and members of the Carbone and Chavez lab for valuable feedback on the research. Authors acknowledge the ENCODE Consortium and Dr. Myers at HudsonAlpha Institute for Biotechnology, who generated the human PU.1 ChIP-seq data. Gibbon PU.1 ChIP-seq was performed by the Epigenetics Consortium at Knight Cardiovascular Institute of Oregon Health and Science University (OHSU). All ChIP-seq libraries were sequenced at the OHSU Massively Parallel Sequencing Shared Resource and the Genomics and Cell Characterization Core Facility at University of Oregon. Data analyses were performed on the Exacloud super computer cluster at OHSU. This work was financially supported by a grant awarded to L.C. from the Leakey foundation. L.C. and R.O.N are also supported by the National Science Foundation (1613856), L.C. and N.A. are supported by the National Human Genome Research Institute (R01HG010333) and L.C. is supported by the NIH/OD P51 OD011092 to the Oregon National Primate Research Center. J.D.F is supported by F32GM125388 and S.R.S is funded by 1R01HG010329. Authors declare no conflict of interest.

## Supplemental Methods

### *In silico* validation of the MELT pipeline

Considering the composite structure of LAVA (Fig. 1A), and the lower quality of the gibbon genome assembly compared to human, selecting the right software for identifying LAVA insertion/deletions (indels) from short-read sequencing data was critical. We selected the MELT software (*1*), which can use information from short-read whole genome sequencing (WGS) alignments, to identify both TE insertions and deletions that are not present in a reference genome. Briefly, MELT identifies TE insertions absent in the reference genome (non-reference) by finding discordant read pairs aligning to the consensus TE sequence and identifies TE deletions based on alignment discrepancies at the known positions of TEs in reference genomes. At last, these predictions are merged across all WGS datasets to fine-tune predictions and genotype (presence/absence) individuals at each TE indel site (*1*).

To test MELT’s (*1*) ability to identify LAVA insertions and deletions (indels) from gibbon WGS data, we simulated 9 mock WGS datasets by introducing LAVA insertions and deletions (indels) in the gibbon genome reference (Nleu3.0) *in silico*. Briefly, we removed all short (<1Mb) and unplaced contigs from Nleu3.0. Next, we generated *in silico* LAVA insertions by randomly inserting sequences of different full-length (1.5-2 Kb) LAVA into Neul3.0 using SVsim (available at https://github.com/GregoryFaust/SVsim). *In silico* deletions of LAVA elements previously annotated in Nleu3.0 were generated with BEDtools (*2*) and custom bash scripts. Using this approach, we generated three modified genome sequences, each containing 100 different insertions and 20 different deletions (mock genomes 1, 2 and 3) and then used wgsim (available at https://github.com/lh3/wgsim) to simulate 100 bp paired-end Illumina reads (fragment length of 250 bp, mutation rate=0.01 and indel fraction=0.01) from each of them (30X coverage, ~421 million read pairs). Simulated WGS datasets were then aligned to Nleu3.0 using bwa (*3*) with default paired-end settings. After using SAMtools (*4*) to binarize and sort alignments, each alignment was down-sampled to 20X and 10X coverage using Picard Tools (available at http://broadinstitute.github.io/picard). We then used MELT v2.1.3 (*1*) with default settings to predict non-reference LAVA insertion and deletion (indel) on each of the 9 simulated WGS datasets.

We first assessed the precision of MELT’s position predictions by comparing predicted vs. true positions of the simulated LAVA indel loci. For both deletions and insertions, the 10X coverage simulation datasets had the lowest percentage of LAVAs predicted precisely at the correct genomic position (58.3 ± 4.7% of insertions, 10 ± 5% of deletions; mean ± stdev; Fig. S1A). This percentage improved by increasing the coverage to 20X (65.7 ± 5.9% of insertions, 13.3 ± 7.6% of deletions; mean ± stdev), but further improvements were minimal after increasing the coverage from 20X to 30X (66 ± 6.1% of insertions and 13.3 ± 7.6% of deletions; mean ± stdev; Fig. S1A). By considering LAVA positions “correct” if they were predicted within 10 bp of their true simulated indel site, the percentage of correct positions increased drastically across all coverages [insertions at 10, 20 and 30X: 92 ± 1.7%, 96 ± 0.0% and 96.7 ± 0.6% (mean ± stdev) and deletions at 10, 20 and 30X: 70 ± 13.2%, 81.7± 7.6% and 81.7± 7.6% (mean ± stdev)]. However, the effect of coverage remained broadly the same, with noticeable improvement obtained by increasing the coverage from 10X to 20X, but not from 20X to 30X (Fig. S1A). Interestingly, increasing the margin of error for predicted position to −/+ 100 bp, or −/+1 Kb of true simulated indel site, did not drastically improve the percentage of correctly predicted LAVA indels (Fig. S1A), suggesting that MELT detected most LAVA within 10 bp from their true indel site.

In order to formally measure specificity (true negative rate) and sensitivity (true positive rate) of our MELT pipeline in identifying LAVAs across the three WGS coverages (10X, 20X and 30X), we allowed for a 10 bp margin of error between the predicted genomic position and the true LAVA indel site and calculated false positive, false negative, and true positive LAVA predictions. Based on these calculations, we observed that the average specificity for predicting LAVA insertions decreased slightly with increase in coverage, but remained high overall (>98%, Fig. S1B, left). Average specificity of prediction LAVA deletions remained 100% across all coverages (Fig. S1B, right). MELT’s average sensitivity in predicting *in silico* simulated LAVA insertions was high across coverages and increased with higher coverage (>92%, Fig. S1B, left). Similarly, average sensitivity for detecting *in silico* simulated LAVA deletions increased from an average 70% in 10X WGS datasets to an average of 81.7% in 20 and 30X coverage simulations (Fig. S1B, right).

### Assessing distribution of LAVA insertions across and within chromosomes

To test whether the number of observed fixed- and poly-LAVA insertions per chromosome deviated significantly from random expectation, we used BEDtools (*2*) to shuffle LAVA insertion sites across the Nleu3.0 genome 1000 times, and each time we counted the number of LAVA insertion loci found in each chromosome. We used these values to measure the mean and standard deviation of LAVA insertions expected per chromosome (Fig. S2A). We then used a two-tailed chi-squared test using the Graphpad tool (https://www.graphpad.com/quickcalcs/contingency1.cfm) to assess whether the overall observed LAVA distribution across chromosomes was significantly different from mean expected distribution. We also determined one-tailed empirical p-values for the deviation from expectation per chromosome, by measuring the proportion of times the shuffled values were more extreme than insertions per chromosome (Fig. S2A).

To test whether LAVA insertions were located closer to each other than expected by random chance (i.e. clustering), we used a permutation approach similar to Kostka et al. (*5*). Briefly, we computed the median of distances between each LAVA insertion and its next nearest LAVA insert site (513.9 Kb median distance for fixed-LAVA, and 530.3 Kb for poly-LAVA). Next, we used BEDtools shuffle (*2*) to shuffle the LAVA insertions within their original chromosome 1,000 times, while avoiding assembly gaps. After each random shuffle, we calculated the median of the shortest distances between LAVA insertion sites (820.3 ± 41.7 Kb for fixed-LAVA, and 794.8 ± 39.5 Kb for poly-LAVA; mean ± stdev of medians). The proportion of the shuffled median distances that were smaller or equal to the true median value was our permutation p-value.

### Assessing distribution of LAVA relative to repeats and genes

In order to test abundance of repeats nearby LAVA, we used the TEanalysis tool [available at https://github.com/4ureliek/TEanalysis, (*6*)] to calculate the percentage of fixed- (n= 1,095) and poly-LAVA (n=1,171) that have inserted within 1 Kb of various repeat families (making sure to exclude the focal LAVA from the 1 Kb window). TEanalysis compares these percentages to those calculated by 1,000 random permutations of LAVA positions, to assess which repeat families are found within 1 Kb of LAVA elements more or less often than expected by random chance (i.e. over-/ and under-represented repeats, respectively). P-values from this analysis are corrected for multiple testing using the Benjamini-Hochberg method (*7*).

In order to examine LAVA’s distribution relative to genes, we first used BEDOPS v2.4.32 (*8*) to calculate the shortest distance between each fixed- and poly-LAVA and gibbon genes. Next, we shuffled fixed- and poly-LAVA 1000 times within their chromosomes (avoiding assembly gaps) using BEDtools (*2*). After each shuffle, we calculated the mean of shortest distances between LAVA and genes (19.5 Kb for fixed-LAVA and 17.6 Kb for poly-LAVA). The proportion of times the mean of the shortest distance between genes and shuffled-LAVA [34.6±2.2 Kb (mean±stdev) for shuffled fixed-LAVA and 33.2 ±2.2 Kb (mean±stdev) for shuffled poly-LAVA] were smaller than that of true LAVA was our empirical permutation p-value.

Lastly, we used the same 1000 shuffled sets of fixed- and poly-LAVA to calculate the expected percent overlap between LAVAs and gene structures (i.e. intergenic, terminator/promoter, exon, UTR and intron). LAVA was considered significantly enriched in a given gene structure (empirical p-value <0.05) if the percent observed overlaps between fixed- or poly-LAVA was higher than >95% of the shuffled datasets. LAVA was considered significantly depleted (empirical p-value <0.05) if the percent observed overlaps between fixed- or poly-LAVA was smaller than >95% of the shuffled datasets.

We used custom R scripts to perform linear regression between LAVA and gene density across chromosomes, where density of LAVA/genes was calculated as (# LAVA/gene in chromosome)/(Mb ungapped chromosome length).

### Establishment of gibbon EBV-transformed lymphoblastoid cell lines

In this study, we used two Epstein Barr Virus (EBV) transformed lymphoblastoid cell lines established in the past (*9, 10*), and using the same method we established cell lines from seven additional NLE gibbons. Briefly, whole blood from gibbons was collected opportunistically in sodium heparin tubes during routine check-ups at zoos or the Gibbon Conservation Center (Santa Clarita, CA). We isolated lymphocytes from the blood using Ficoll-Paque PLUS (GE Healthcare). Next, we transformed 3-9×10^6^ lymphocytes with EBV from the marmoset cell line B95-8 (ATCC CRL-1612), using a standard protocol. Briefly, we incubated the cells with EBV for 2 hours at 37°C then diluted with RPMI-1640 (Corning cellgro) supplemented with 10% FBS (Hyclone), 1X MEM Non-essential Amino Acids Solution (Corning cellgro), 1mM Sodium pyruvate (Corning cellgro) 1% Pen-Strep (Corning cellgro) and 2mM L-glutamine (Hyclone). Lastly, we allowed the cells to grow undisturbed for 10-12 days. Once signs of transformation were observed, we fed the cells with the same supplemented RPMI-1640.

### Histone chromatin immunoprecipitation (ChIP), library preparation and sequencing

To perform each ChIP assay, we fixed 5×10^6^ cells with 1% formaldehyde for 5min on ice and then quenched fixation by adding glycine to a final concentration of 0.1M. After washing the cells twice with cold 1X PBS, we lysed the fixed cells at a concentration of 3×10^6^ cells/100 ul in lysis buffer [0.1% SDS, 0.5% Triton X-100, 20mM Tris-HCl pH=8.0, 150mM NaCl, 1x Proteinase inhibitor (Roche)] for 5min on ice. Using the Bioruptor Pico sonicator (Diagenode), we sheared lysates in 1.5mL tubes with 7 cycles (30sec on/off). We spun the sonicated lysates at max speed for 10min at 4°C to remove cell debris and then rotated them for 2hr at 4°C with 20 ul ChIP-grade Protein A/G Magnetic Beads (Pierce) to preclear the lysate. From each chromatin preparation, we set aside a 1% volume aliquot as chromatin input, and divided the remaining chromatin into equal volumes for each of the five histone mark ChIP reactions. We diluted the reactions to 1.6×10^6^ cells/100 ul with lysis buffer and then added the corresponding antibodies in the following amounts: 2 ug H3K4me1 (ab8895, Abcam), 1 ug H3K4me3 (ab8580, Abcam), 1 ug H3K27ac (ab4729, Abcam), 2 ug H3K27me3 (39155, Active Motif) and 2 ug H3K9me3 (ab9263, Abcam), and let reactions rotate overnight at 4°C. Next, we added 20 ul Pierce ChIP-grade Protein A/G Magnetic Beads (Pierce) and let samples rotate for 2hrs at 4°C. We then washed the beads consecutively (rotating at 4°C) with TBST buffer (1X Tris-Buffered Saline, 0.1% Tween; 3 times for 15min), lysis buffer (once for 1hr), 1X TE pH=8 (once for 1hr, then once for 10min). Following washes, we eluted chromatin in 250 ul fresh elution buffer (1% SDS, 0.1M NaHCO_3_). We incubated the ChIP and input samples overnight at 65°C in the presence of NaCl to reverse crosslinks and then digested them consecutively with 8 ug RNAse A (30 minutes at 37°C) and 80 ug Proteinase K (2 hour at 55°C). Lastly, we purified the samples using traditional phenol:chloroform extraction and ethanol precipitation. We quantified all samples using the Qubit dsDNA High Sensitivity kit (Thermo Fisher Scientific).

All sequencing libraries were generated using the NEBNext Ultra II DNA Library Prep Kit for Illumina (New England Biolabs) from 1-5 ng of starting material and without size selection. The Qubit dsDNA High Sensitivity kit (Thermo Fisher Scientific) and Agilent Bioanalyzer 2100 were used to QC the libraries. Histone ChIP libraries were sequenced on the SE 75 bp Illumina NextSeq and the SE 100 bp Illumina HiSeq 2500 platform at the Massively Parallel Sequencing Shared Resource (MPSSR) at Oregon Health Science University. Details on sequencing strategy (i.e. single-end vs. paired-end), read counts and alignment statistics for each ChIP library can be found in Table S4.

### PU.1 ChIP-seq, analysis and assessing binding to LAVA and SVA

ChIP against PU.1 was carried out in the Epigenetics Consortium of the Knight Cardiovascular Institute at Oregon Health and Science University similarly to the histone ChIP procedure described above, with the following differences: 10×10^6^ fixed cells were used per ChIP assay, 6 ul of the PU.1 antibody (MA5-15064, ThermoFisher Scientific) was used for pull-down and fixed cells were only sonicated for 5 cycles (30sec on/off). Library preparation was performed as described above. PU.1 ChIP-seq libraries were sequenced on the PE 2×100 Illumina HiSeq 4000 platform at the Genomics and Cell Characterization Core Facility (GC3F) at the University of Oregon.

Raw reads from in-house gibbon and public human PU.1 ChIP-seq (GEO accession no.: GSM803531 and GSM803398) were QC’d using FastQC (*11*) and trimmed with Trimmomatic (*12*). We first used bowtie2 (*8*) with default settings to align processed reads from gibbon and human ChIP-seq replicates to Nleu3.0 and Hg38, respectively. RepEnrich2 (*13*) and edgeR (*14*) were next used to calculate number of aligned reads to each repeat family and to test enrichment of repeat families in PU.1 ChIP-seq data compared to input, per developers’ instructions (*13*). Briefly, we removed all low-complexity and simple-repeat annotations from the RepeatMasker annotations of each genome. The modified RepeatMasker annotations were used by RepEnrich2 (*13*) along with the ChIP and input alignments to calculate the amount of unique and multi-mapping read alignment to repeats in each dataset. Next, edgeR (*14*) was used to normalize read counts based on library size (counts per million, CPM) and to test significant enrichment of all repeat families in PU.1 samples relative to input (FDR <0.05). While the human and gibbon datasets had different read lengths (SE 1X36 bp vs. PE 2×100 bp; Table S4), we do not expect this to impact our results, since repeat enrichment is determined by comparing alignment across ChIP and input datasets sequenced with the same read lengths.

To characterize which regions of LAVA are bound by PU.1, we next created PU.1 ChIP-seq meta-summits, as described before (*15*). Briefly, we first created the consensus LAVA sequence by aligning 50 of the longest full-length LAVA elements present in the Nleu3.0 genome using MUSCLE (*16*). The longest 1% of LAVA elements, which often contain mis-annotations or recombination products, were not used. The resulting alignment was then used to generate a consensus (stripping columns that were >95% gaps) as previously described (*15*). This consensus was then annotated using TRF [Tandem Repeat Finder, (*17*)] to identify the VNTR region. To identify the *Alu*Sz6 and L1ME5 regions, we used BLAT (*18*) to map portions of the consensus back to the genome and identified overlap with genomic *Alu*Sz6 and L1ME5 regions. We then used alignments generated as parts of the RepEnrich2 analysis described above, to identify PU.1 peaks using MACS2 (*19*), while allowing multi-mappers. We used BEDtools (*2*) to identify gibbon PU.1 summits within reference-present LAVA. We combined LAVA-overlapping summits from two biological replicates and extended them by 25 bp in either direction. We then used blat to align and pileup these summits to the LAVA consensus. In this approach we use each instance of overlap between a PU.1 peak and a LAVA element as a “replicate”, therefore the pileup we create along the consensus represents the combination of overlaps across many genomic replicates. Summits of these pileups (i.e. “meta-summits”) were determined by running MACS2 (*19*) using *--keep-dup all --nomodel --call-summits --extsize 50* parameters. All data is available as a genome browser session at: http://genome.ucsc.edu/s/jdf2001/LAVA_paper. PU.1 meta-summit analysis in human were done similarly to gibbon, using alignments from RepEnrich2 alignments (described above) and publicly available Human SVA_A consensus from the UCSC repeat browser (*15*).

Lastly, to identify specific LAVA elements binding to PU.1, we aligned gibbon ChIP-seq data using BWA with default settings (*3*), removed low quality and multi-mapping read alignments using samtools (*4*) and used MACS2 (*19*) to predict significant peaks (q<0.01) relative to appropriate input. We determined the correlation between the biological replicates using deepTools2 (*20*). Since the replicates displayed high correlation (Pearson correlation coefficient=0.82), we pooled the gibbon PU.1 peaks (Table S4) across the two replicates and merged overlapping peaks using BEDtools (*2*). This resulted in a total of 13,920 unique ChIP-seq peaks, which were significantly enriched for the consensus PU.1 recognition motif (Fig. S5C) and co-occurred with active histone marks (Fig. S5D). We intersected PU.1 peaks and reference-present LAVA inserts using BEDtools (*2*) by requiring the PU.1 peak summit to fall within LAVA.

### RNA-seq data collection and normalization

Total RNA was extracted from the nine EBV-transformed lymphoblastoid cell lines (LCLs) we established for NLE individuals using the RNeasy Mini kit (Qiagen), following manufacturer’s instructions. RNA integrity scores were assessed on the Bioanalyzer with the RNA 6000 Pico kit (Agilent Genomics). RNA-seq libraries were generated with the TruSeq kit and sequenced to obtain ~40 million SE 100 bp reads/sample at the OHSU Massively Parallel Sequencing Shared Resource. Reads were trimmed with Trimmomatic (*12*) and aligned to Nleu3.0 using STAR (*21*). We counted high-quality uniquely mapping reads (MAPQ>25) aligning to each gene and then normalized our gene counts in multiple steps. First, we used edgeR (*14*) to remove genes for which fewer than two individuals had >0.5 read counts per million (CPM). Next, we used the Conditional Quantile Normalization package [CQN (*22*)] to normalize gene counts based on gene length, GC content and RNA-seq library size. The normalized read counts were regularized log (rlog) transformed using DEseq2 (*23*) to stabilize the variance, and lastly, genes located on unassembled contigs in the genome were excluded.

### Expression of genes near LAVA vs. the rest of the genome

We used custom R scripts and a two-tailed permutation approach to compare median of expression of the nearest gene within 3 Kb of genic fixed (n=448) and polymorphic LAVA (n=478), genes elsewhere in the genome. Briefly, for each gene near fixed- or poly-LAVA insertion loci, we calculated the median normalized expression among our 9 NLE individuals, and calculated the median of all these median values for fixed- and poly-LAVA, separately. Then, we randomly selected the same number of genes across the genome (excluding those nearby LAVA) 10,000 times, for comparisons with fixed- (n=448) and poly-LAVA (n=478). After each random selection, we calculated the median of medians of selected genes. Our empirical p-value was the proportion of times (out of 10,000) that the median of medians of randomly selected genes exceeded that of the genes near fixed- and poly-LAVA.

### Allele frequency spectrum estimation and summary statistic estimation

We used ANGSD (*24*) to estimate a folded allele frequency spectrum (AFS) for 10 *Nomascus leucogenys* (NLE) genomes, excluding the individual used to construct the reference genome. This method uses genotype likelihoods to estimate population genetic parameters, rather than downstream genotype calls, and thus can better account for lower or variable coverage across samples (four samples had coverage <20x). Individual genotype likelihoods were calculated using the GATK approach (-GL 2) and assuming Hardy-Weinberg equilibrium (-doSaf 1). In addition to estimating AFS for the whole genome, we also estimated AFS separately for two sets of putatively neutral regions. The first was for ~12,000 putative 1 Kb non-genic loci identified in Veeramah et al. (*25*) (total length =124,073,560 bp), and the second was for ~34,000 putative 1 Kb neutral loci lifted over from Hg18 from Gronau et al. (*26*) (total length = 35,587,610 bp).

Genome-wide AFS was used by ANGSD as the basis for estimating genome-wide summary statistics for the same set of 10 NLE individuals. Summary statistics (e.g. ΘW, Θπ and Tajima’s D (*27*) were calculated in windows of 10 Kb, and a stepping size of 1 bp. From this point forward, when describing the Tajima’s D value of a LAVA element (both fixed- and poly-LAVA), we are referring to the 20 Kb flanking its insert site (i.e. 10 Kb upstream and 10 Kb downstream). When plotting regional Tajima’s D, a nonparametric lowess function was used to smooth across the sliding windows with a delta equal to the range of values within the region of 0.01, and using 10% of the data to calculate each y value. Within any 10 Kb window for which we estimate a summary statistic, some set of positions may not be “callable” by ANGSD (*24*). Callable sites refer to positions in the genome where ANGSD could successfully estimate genotype likelihoods, and thus summary statistics. Failure to do so may be due to lack of coverage or an ‘N’ in the ancestral sequence. Thus, any 10 Kb locus may have between 0 and 10,000 callable sites.

### Coverage-filtering of LAVA elements

In order to limit the potential impact of copy number variation or segmental duplications in our analysis due to incorrectly called SNPs, we calculated the average per site coverage for the 20 Kb (10 Kb either side) surrounding each LAVA element across all 10 NLE individuals. We used average coverage across chr2 (258x) as a guide to identify putatively “clean” LAVA elements, and only retained LAVA elements with total coverage between 200 and 300x for downstream selection analysis (fixed LAVA elements n=734/1095, polymorphic LAVA elements n=808/1171). This criterion excluded all LAVA elements on chrX. Of this subset of 734±808 loci, we also examined mean WGS coverage of their corresponding chromosome in individuals. Of the 1,542 LAVA elements, only 13 had coverage that was more extreme than double or half their mean chromosomal coverage. Only one poly-LAVA element (chr2:112689756-112689757) showed coverage more than double the chromosomal mean in three individuals, likely representing the only true CNV still remaining in our data [all other loci showed marginally extreme coverage values that might be expected by chance given the large number of comparisons (n=15,420)].

### Assessment of Tajima’s D significance under empirical framework

As described above, we used ANGSD (*24*) to estimate various population genetic summary statistics across the genome in 10 Kb windows sliding every 1 bp using ten NLE whole genomes. In particular, we focused on Tajima’s D (*27*), a statistic that measures the difference between two estimates of θ. Significantly negative Tajima’s D values indicate an excess of singletons compared to neutrality, compatible with a scenario of either a recent selective sweep or background selection. Significantly positive Tajima’s D values indicate an excess of intermediate frequency alleles compatible with a scenario of balancing selection. It should be noted that negative and positive Tajima’s D values can also result from population expansions and contractions (i.e. demographic events), respectively. However, these will affect the whole genome, rather than any single locus.

We utilized two major frameworks (empirical vs. simulation) to assess significance of the negative Tajima’s D values at individual LAVA elements, indicative of positive or purifying selection. In the empirical framework, we compared the Tajima’s D upstream and downstream of LAVA elements to all other loci from across the genome that were considered callable, randomly sampling 10,000 loci from this set to avoid correlated windows. This allowed us to generate empirical p-values for each individual LAVA element (one for the upstream section and one for the downstream section). We considered a LAVA element to be potentially under selection only if p<0.05, for both the upstream and downstream region. To be analyzed, loci needed to be callable at least 25% of sites (i.e. ≥2.5 Kb within the 10 Kb window).

### Assessment of Tajima’s D significance under simulation framework

In order to generate p-values based on neutrality rather than just being outliers in the genome (i.e. simulation versus empirical p-values), we used ∂a∂i (*28*) to fit a demographic model to the AFS derived from the two different sets of putatively neutral loci (*25, 26*). We tested the likelihood of both a standard neutral model (snm) and a one-step size change (2-epoch) model. For the latter, the free parameters were the relative size change from the ancestral population effective population size (nu) and the time of this size change in coalescent units (t). Likelihoods between the simulated and observed data were assessed using the scaled multinomial method, with θ estimated after fitting the best model. Likelihoods for different combinations of nu and t were first examined along a two-dimensional grid with nu ranging from log(nu)=-3 to log(nu)=3 in steps of 0.1, and t ranging from 0 to 2.0. The best estimate from this grid was then used as a starting point to fit the data using the Broyden-Fletcher-Goldfarb-Shanno (BFGS) optimizer. The grid sizes for extrapolating the approximate solution to the partial differential equation was 40, 50, 60.

A 2-epoch model with a population expansion in the recent past provided a much better fit to the data compared to a simple standard neutral model (Fig S8). Estimates of the relative size change (nu) and time of this size change (t) were very similar for both data sets (Table S5) and are suggestive of a ~four-fold increase in population size ~50,000 generations ago assuming a per generation mutation rate per site of 1×10^−8^. However, we note that in this study were not interested in the exact demographic model, only that it can suitably replicate the observed AFS. In this case, simulating under the best fit 2-epoch model gave an expected Tajima’s D that matched, or was very close to, the observed data (−0.816 vs. −0.813, and −0.821 vs. −0.821 for the Veeramah et al. (*25*) and Gronau et al. (*26*) data, respectively).

Given the model inferred above, a set of 10,000 simulations were performed for each LAVA element to represent the expected distribution of Tajima’s D under neutrality, with these simulations reflecting the same size and missingness level as the LAVA element being analyzed. We utilized both a standard rate of 1 × 10^−8^ per site per generation recombination rate for each locus, as well as a recombination rate of 0. This latter is an extreme case, but we included it as a conservative measure, since Tajima’s D is known to be decreased due to smaller recombination rates (*29*). Unlike the empirical tests above, a single Tajima’s D value was calculated for the upstream and downstream region for both the LAVA element and simulations, and p-values estimated by comparing the former to the distribution of the latter.

The fixed LAVA elements identified as significant using the empirical threshold (both upstream and downstream loci p<0.05) were also found to be significantly negative (p<0.05) using neutral simulations with no recombination (the most conservative case, see methods above), except for two loci which were close to significance (0.053 and 0.51). Twelve additional LAVA elements were found to be significant using the simulation framework but not the empirical analysis, though in all these cases either the upstream or downstream region was significant for the latter. Similarly, the eleven polymorphic LAVA elements found to be significant in the empirical analysis were found to also be significant using the simulation framework. Eight additional polymorphic LAVA elements were found to be significant using the simulation framework only, with either the upstream or downstream loci being significant in the empirical framework (Table S5).

### Tajima’s D analysis of orthologous regions in human

To investigate signatures of selection in human, we first downloaded alignments for 20 randomly sampled individuals from the Yoruba population in the 1000 Genomes Project (*30*) and used them to estimate the folded AFS and calculate genome-wide windowed Tajima’s D via the same ANGSD framework as described for gibbons above.

We then used the multiz100way alignment to find orthologous human regions Hg19) for the 10 Kb lying either side of each LAVA element in Nleu3.0. Of the 734 fixed LAVA elements, human orthologues for both the entire upstream and downstream sequence were found for 672. Of these, 454 were used in further analysis based on the criteria that the size of the human homologue was within 20% of the original gibbon sequence (i.e. between 8kb-12kb) and both upstream and downstream loci were found on the same human chromosome. Similarly, for the 808 polymorphic loci, human orthologues were found for 752, and 581 were used in further analysis. These regions were then assessed for significance of Tajima’s D in the Yoruban individuals using the same criteria utilized in the empirical framework for gibbons.

Amongst the19 fixed LAVA elements with significantly negative Tajima’s D in gibbons, we identified orthologous sequences (via multiple alignments) for 14 elements, but none showed signals of significant Tajima’s D in humsn. For the 8 out of 11 significantly negative polymorphic LAVA elements for which we could find human orthologous sequence, one had significant Tajima’s D in the downstream flanking window. Thus, it is possible that LAVA elements we observe with significantly negative Tajima’s D in gibbons underwent a selective sweep, or experience background selection that is unique to the gibbon lineage (in either case is suggests some functional role for these LAVA elements or the regions directly surrounding them).

### Calculating up/downstream combined Tajima’s D values for co-opted LAVA

We estimated a single “combined Tajima’s D” value for the 10 Kb upstream and downstream windows flanking each of the two putative co-opted LAVA elements for visualization in Fig. 6B. First, we used ANGSD (*24*) to estimate the AFS for the upstream and downstream portions separately, before combining these into a single AFS and estimating Tajima’s D. Next, for each LAVA element we identified 10,000 random pairs of 10 Kb loci that were separated by the same distance as the focal LAVA element and were at least 75% callable. For each random window pair, we calculated a single Tajima’s D using the same ANGDS protocol described before. It should be noted that this analysis, which were specifically carried out for the two putative co-opted LAVA loci, differs from the empirical p-values reported in Table 5S for all the LAVA elements. Adopting this approach to all LAVA was not easily feasible, as calculating the combined Tajima’s D values for two interspersed loci via ANGSD is extremely computationally intensive.

### Validating significance of association between fixed-LAVA and gene ontology terms

To ensure that the significant enrichment of fixed LAVA elements near DNA repair gene ontology (GO) terms was not a result of biases or errors in the gibbon gene annotations, we used two separate permutation approaches to calculate empirical p-values for the significant GO terms identified by Enrichr (*31*). First, we recorded the number of genes found nearby LAVA for each of the significant GO terms. In the first permutation approach, we randomly shuffled the positions of all fixed-LAVAs across the whole genome and recorded the genes within 3 Kb of the shuffled LAVA positions. In the second permutation approach, we randomly shuffled positions of LAVAs that were within 3 Kb of genes, while restricting their new random positions to be <3 Kb of another gene. This latter approach is more conservative and indirectly accounts for effects of gene length (i.e. longer genes are more likely to randomly receive LAVA insertions) and LAVA’s overall tendency to be inserted nearby genes. For both approaches we then identified genes nearby the shuffled LAVA and used Enrichr (*31*) to obtain the GO terms associated with these genes. We repeated each shuffling process 1000 times, and each time recorded the total number of genes associated with each GO term of interest. We then obtained an empirical p-value by comparing the true gene counts for each significant GO term to the null distribution of GO term counts. Using these two permutation tests, we never randomly generated as much (or more) enrichment as we had found with the true LAVA positions for any of the significant GO terms, suggesting that all the GO term enrichments identified by Enrichr were still significant after accounting for imperfections in the gibbon gene annotations (empirical p-value <0.001).

## Supplemental Figures

**Figure S1.**
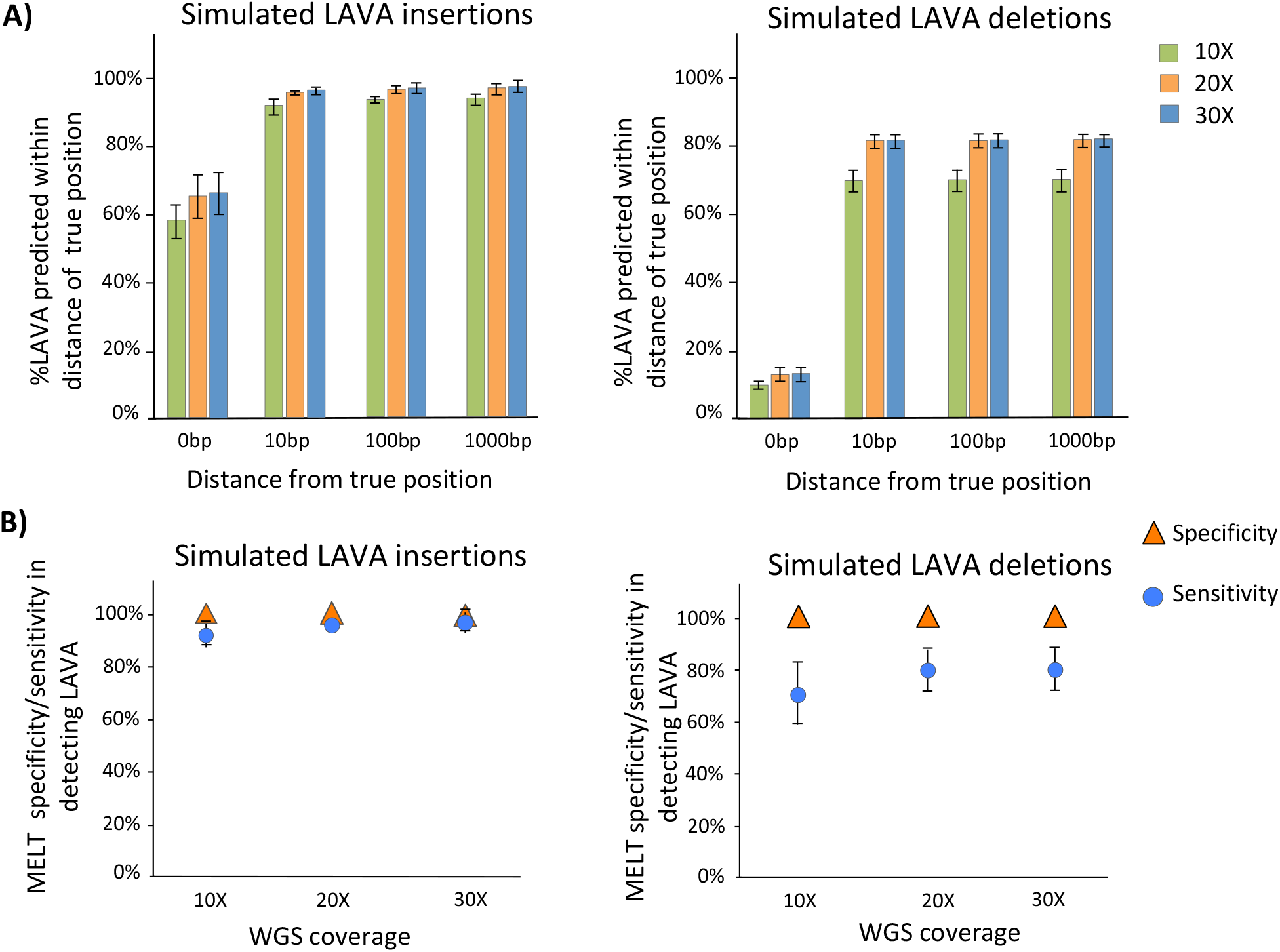
MELT predicts LAVA indels with high precision and accuracy. **A)** Percent *in silico* LAVA insertions (*left*) and deletions (*right*) identified within various distances of their true positions are demonstrated. Bars represent mean of three mock WGS datasets, and error bars represent standard errors. **B)** Percent specificity and sensitivity of LAVA insertion (*left*) and deletion (*right*) predictions by MELT across three coverages. Data points represent mean values across the three mock WGS datasets and error bars represent standard errors.

**Figure S2.**
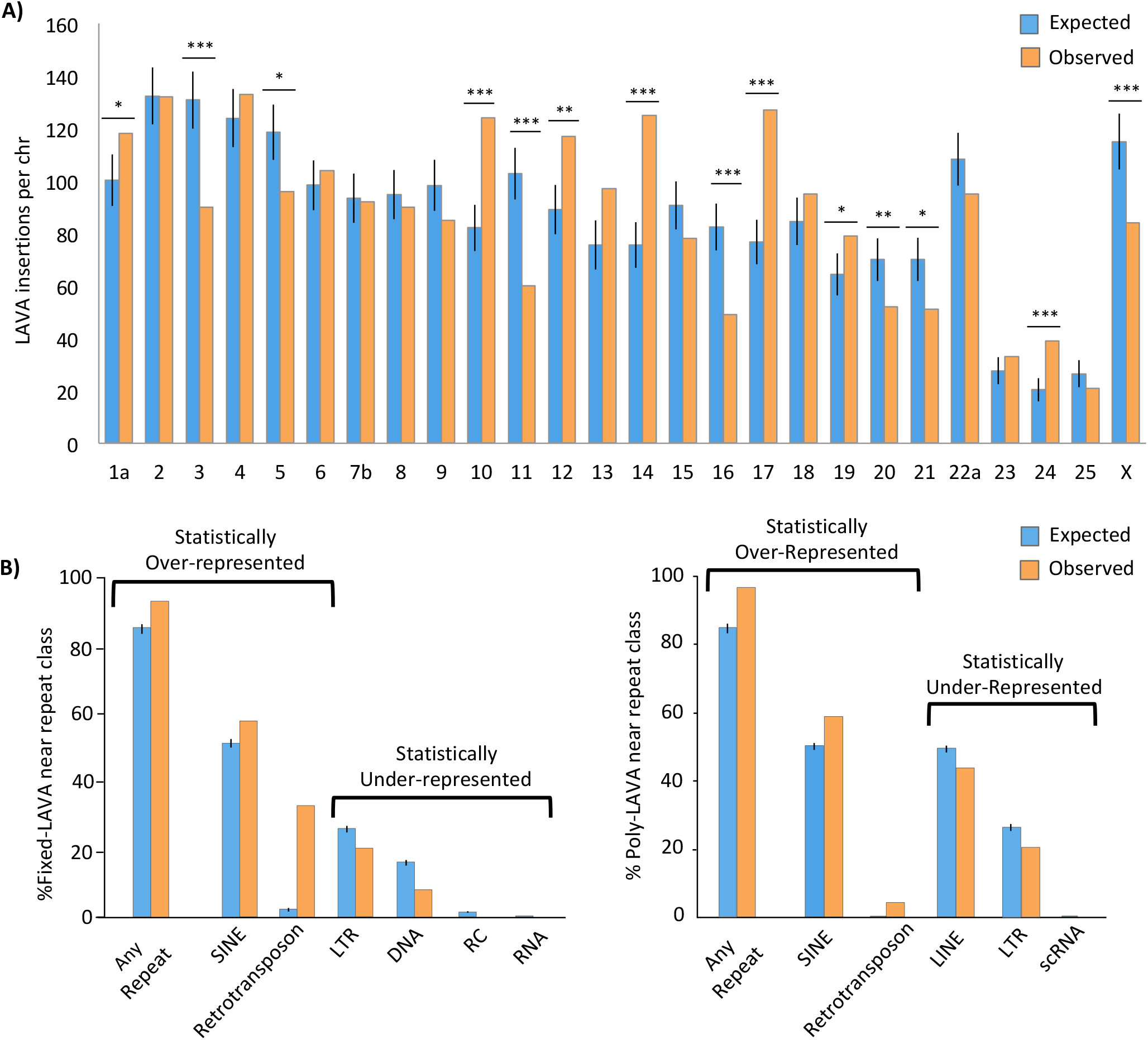
Distribution of LAVA across Nleu3.0. **A)** The average expected number of LAVA insertions per chromosome (based on 1000 shuffles) is plotted against the observed number of LAVA insertions per chromosome. *p<0.05, **p<0.005, ***p<0.0005. **B)** Observed and expected percentages of fixed- (*left*) and poly-LAVA (*right*) LAVA that are within 1 Kb of repeat families are shown. Only repeat families that are significantly over- and under-represented (two-tailed permutation q-value <0.05) within 1 Kb of LAVA are shown. Statistically over (under)-represented means the observed value was significantly larger (smaller) than random expectation. Error bars represent standard error.

**Figure S3.**
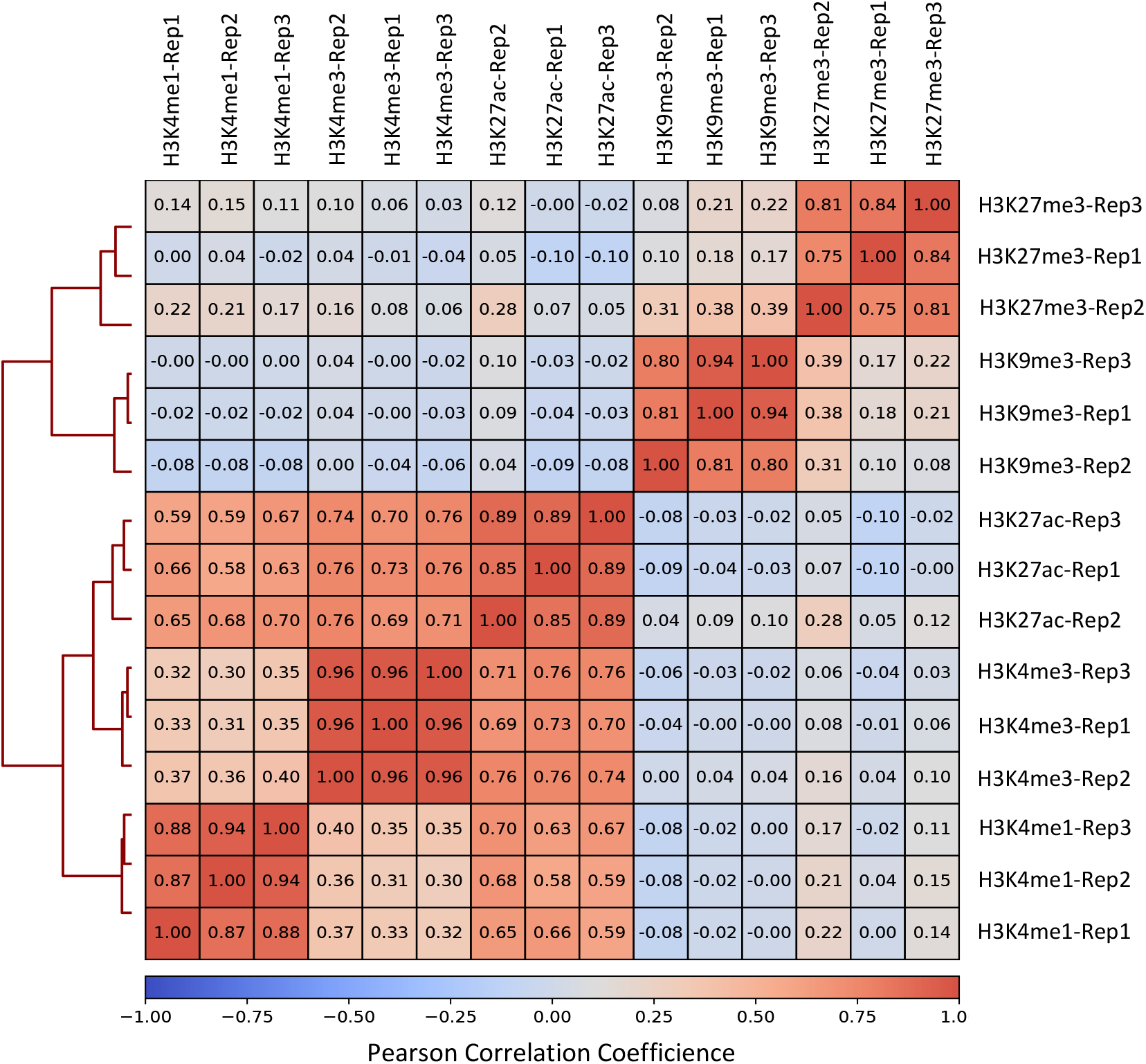
Histone ChIP-seq signals correlate across replicates. A heatmap demonstrates correlation between all histone ChIP-seq replicates. Cell shades reflect Pearson Correlation Coefficient for each pair-wise comparison, with the exact values shown within each cell.

**Figure S4.**
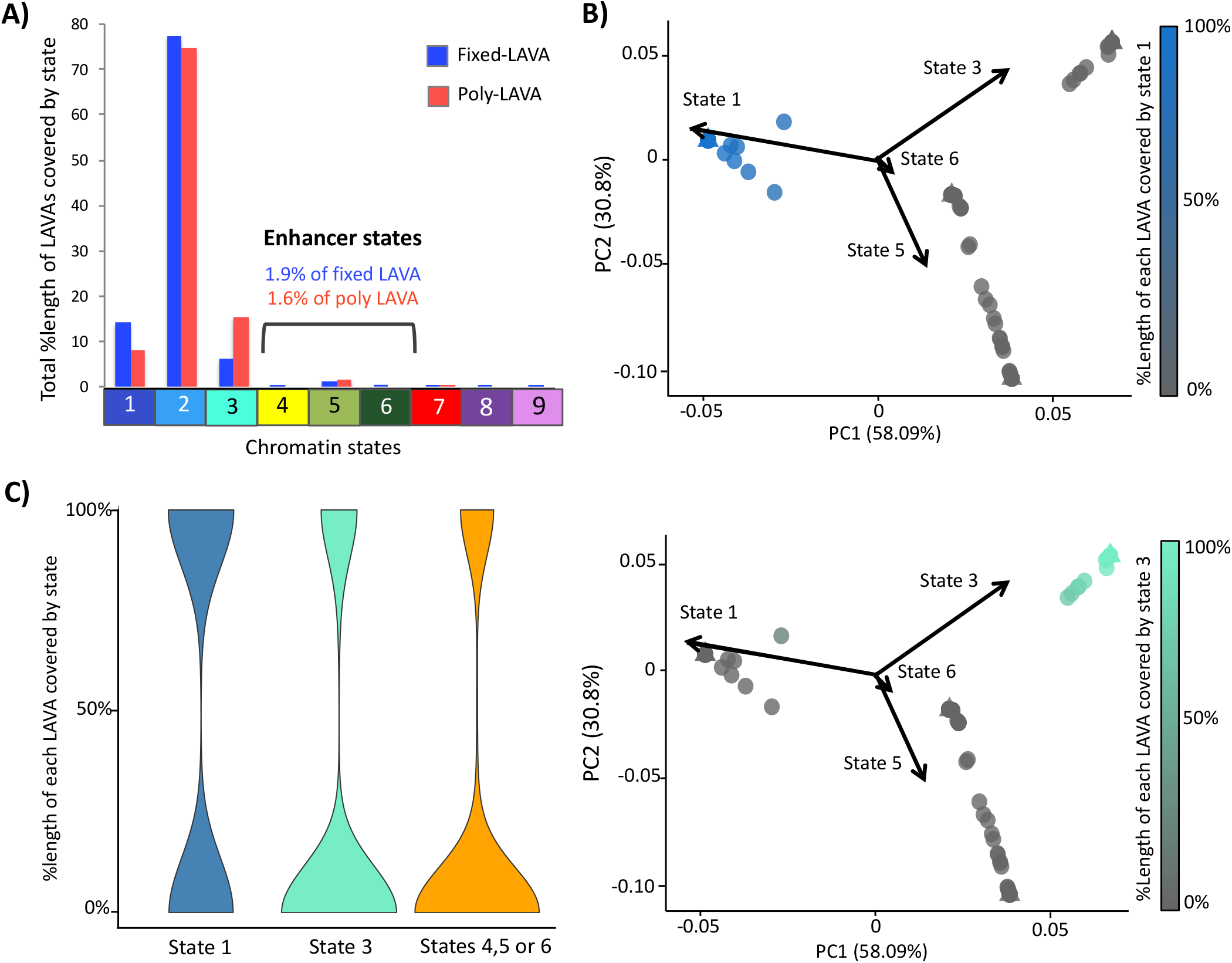
Mappable LAVA can be classified into at least three epigenetically distinct groups. **A)** Percent length of total LAVA sequences that overlap each chromatin state is shown for both fixed (blue) and poly-LAVA (red). Chromatin state and colors match those in Fig. 3A. **B)** PCA biplots group mappable LAVA in three broad clusters. LAVA elements are color coded based on percentage of their mappable length that overlaps state 1 (Constitutive Heterochromatin, *top*) or state 3 (Polycomb Heterochromatin, *bottom*) after excluding regions overlapping state 2 (Low Signal/Mappability). **C)** Violin plots demonstrate density of %overlap between reference-present LAVA elements and Constitutive Heterochromatin State (State 1), Polycomb-Repressed Heterochromatin (State 3) or Enhancer States (States 4, 5 or 6).

**Figure S5.**
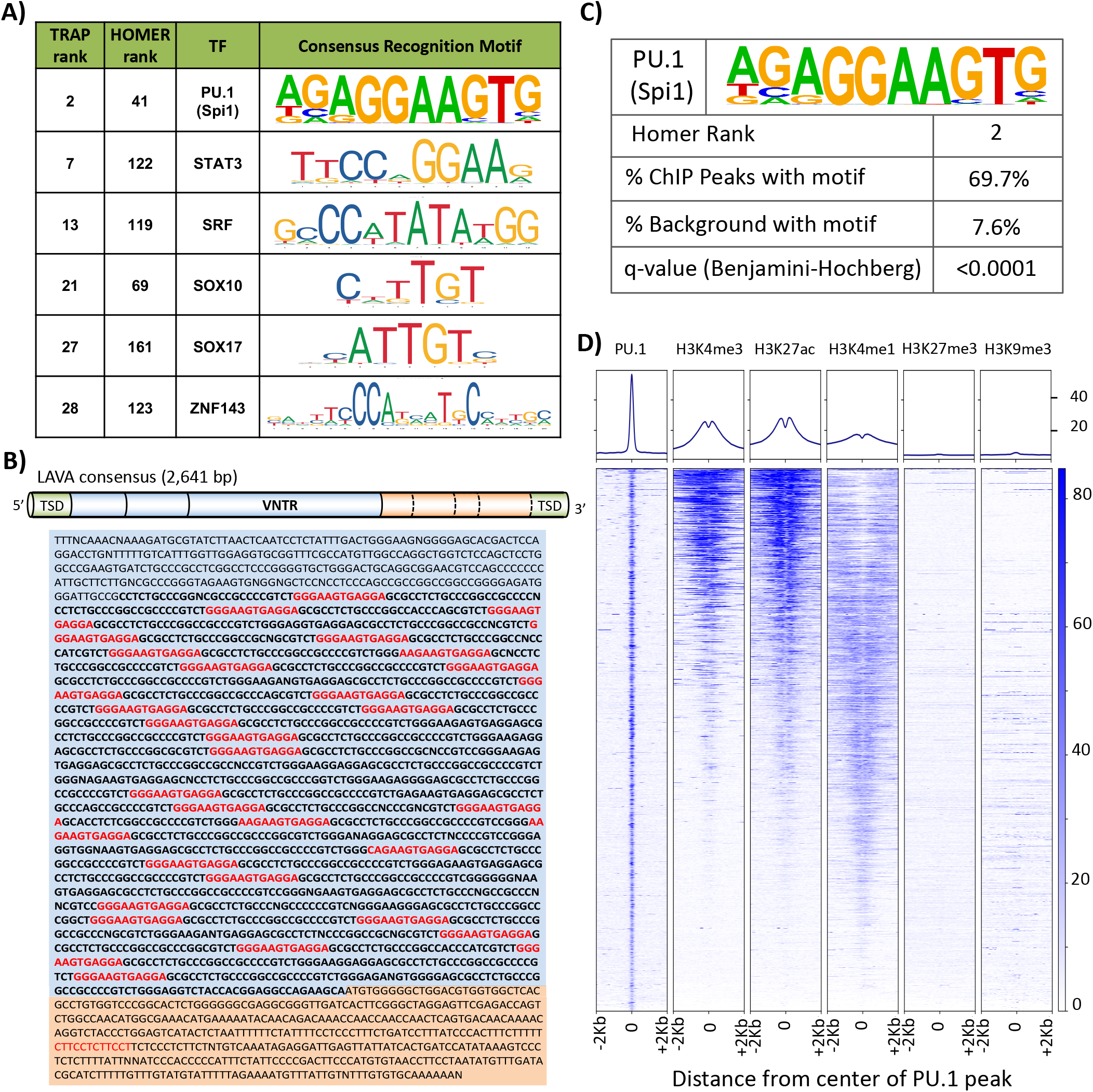
Enrichment of PU.1 recognition motif in LAVA consensus sequence and ChIP output. **A)** Six transcription factor recognition motifs were significantly enriched in LAVA sequences (q<0.05) using both Homer and TRAP pipelines. **B)** Predicted PU.1 binding motifs within consensus LAVA sequence are marked in red. Regions shared with SVA are shaded light blue, and VNTR sequences appear in bold font. **C)** Gibbon PU.1 ChIP-seq peaks are enriched in the consensus PU.1 recognition site. D) Active, but not silencing, histone marks are enriched in 4 Kb windows centered at PU.1 peaks.

**Figure S6.**
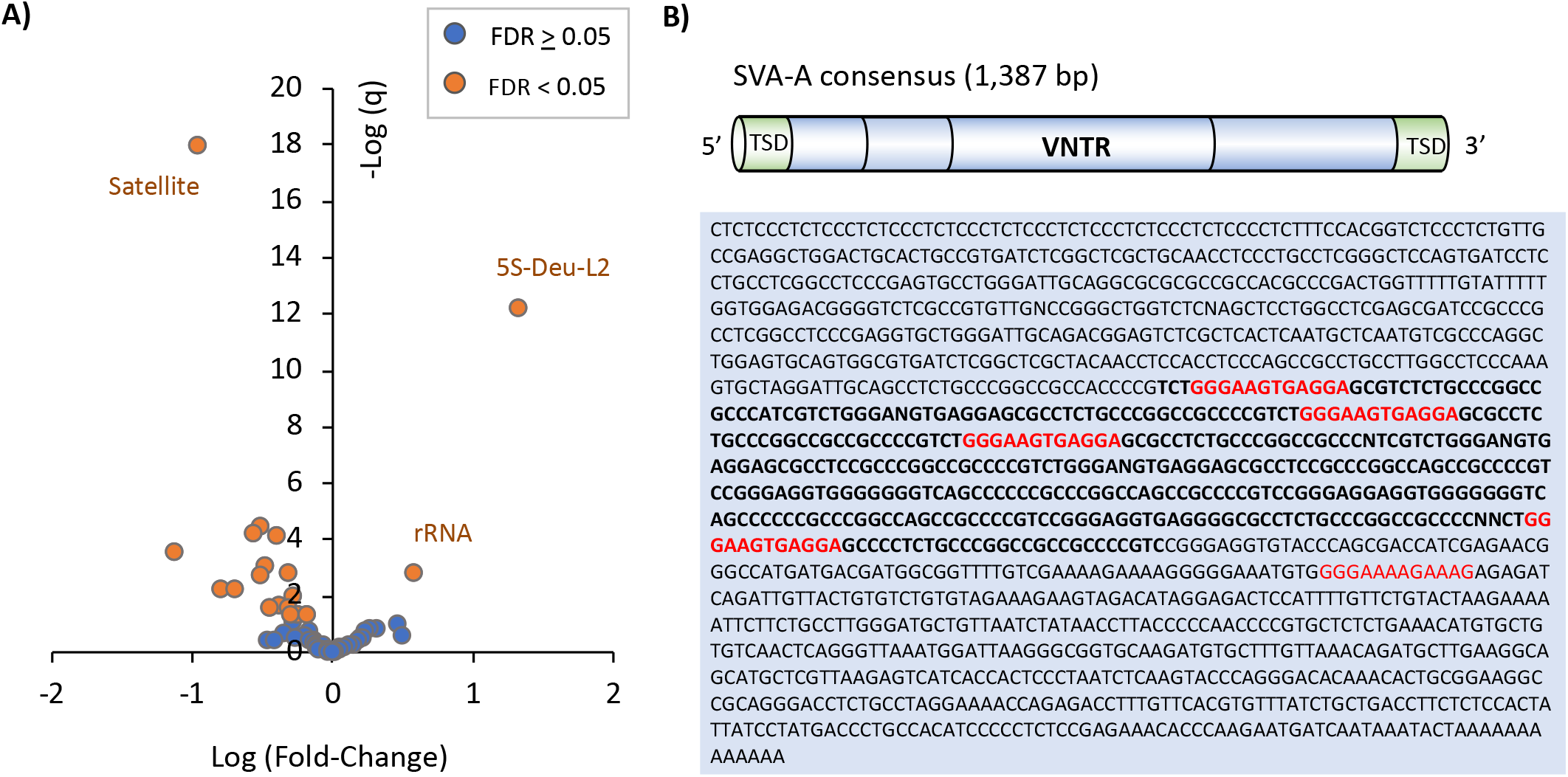
PU.1 does not appear to bind SVA in human. **A)** Volcano plot shows repeat families enriched in public human PU.1 ChIP-seq data. **B)** Predicted PU.1 binding motifs within the SVA_A consensus are marked in red and VNTR sequences appear in bold font.

**Figure S7.**
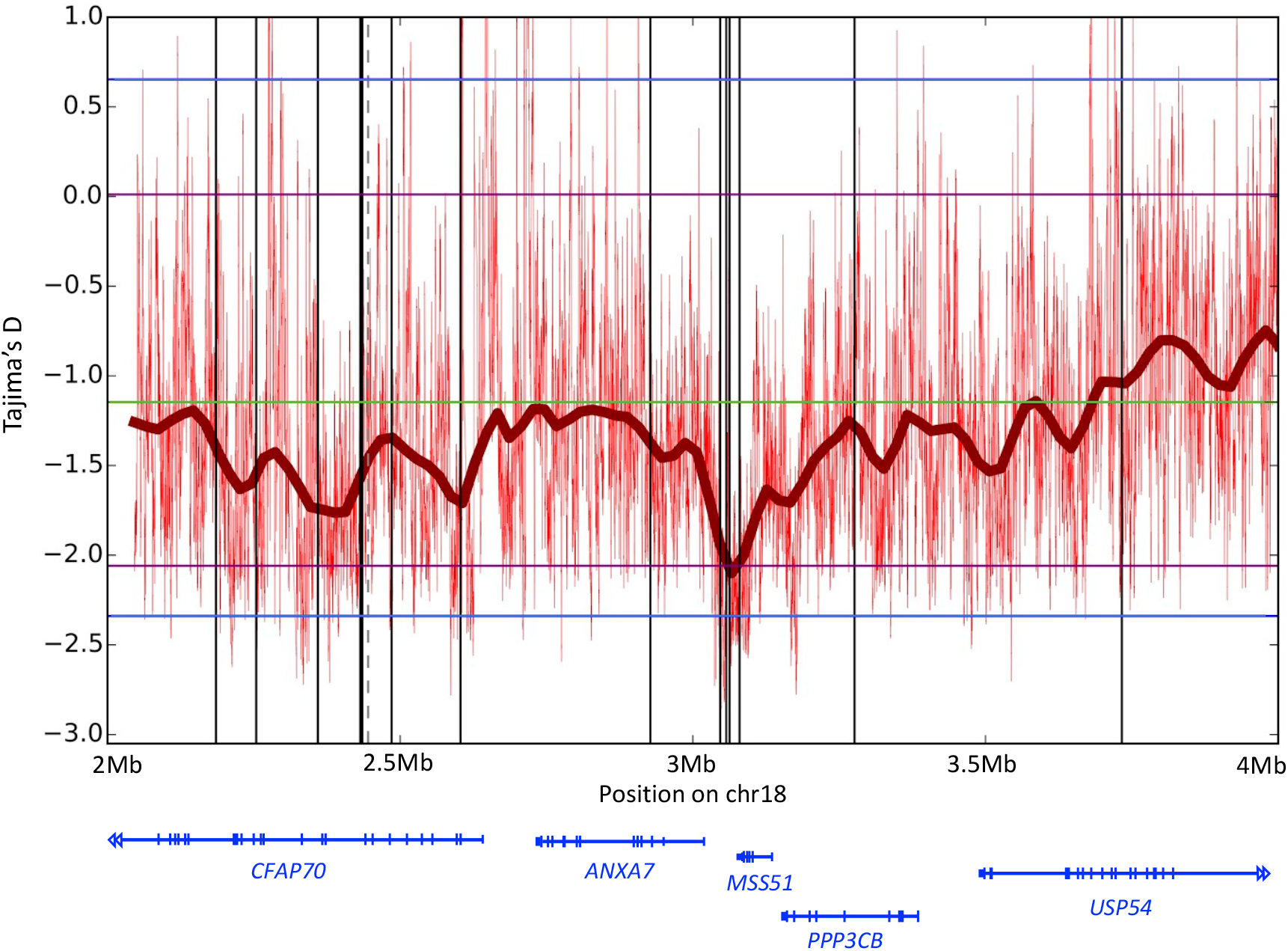
A cluster of four fixed LAVA elements display a strong Tajima’s D dip. Tajima’s D values and genes are shown in a 20 Mb window on gibbon chromosome 18, centered at a cluster of four fixed-LAVA. Red vertical lines show Tajima’s D value of individual 10 Kb windows with 1 bp sliding steps, and thick dark red line is lowess smoothing curve of these windows. Solid and dashed vertical lines mark positions of fixed- and poly-LAVA in this region, respectively. Blue horizontal lines are the 1% and 99% percentile of Tajima’s D across the whole genome, green is the mean across the genome. Tajima’s D at the focal cluster of LAVA is −2.38 (upstream) and −2.39 (downstream). The 99th percentile is −2.34.

**Figure S8.**
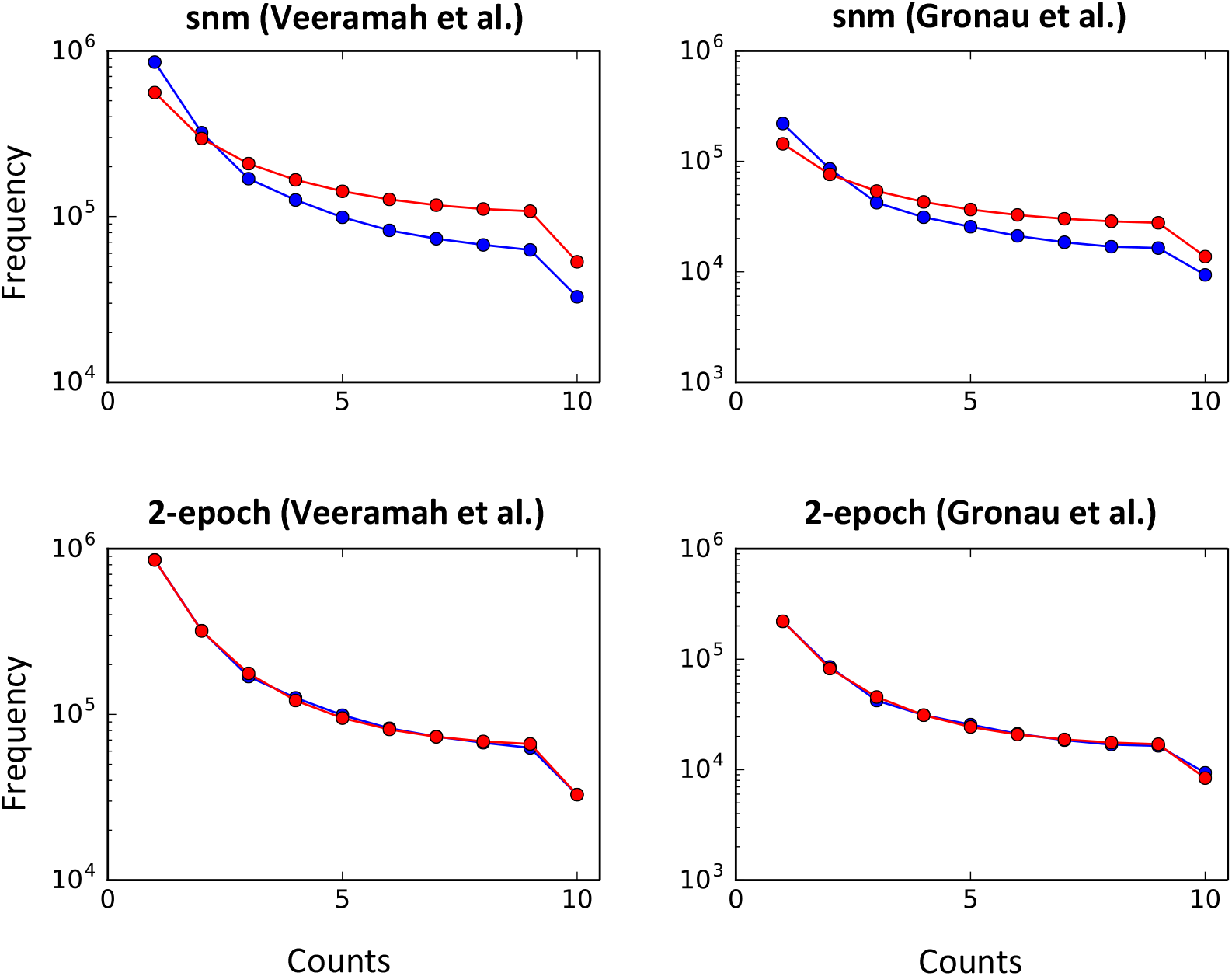
Comparison of standard neutral and best-fit 2-epoch models. Standard neutral model (snm) vs best-fit 2-epoch model from ∂a∂i against folded AFS based on non-genic loci from Veeramah et al. *(25)* (A vs B) and Gronau et al. *(26)* (C vs D). Blue points and lines represent the AFS of the observed data, red points and lines represent the snm and best-fit 2-epoch model. ‘Counts’ refers to number of observations of the minor allele at a particular site among the 10 sampled diploid individuals, ‘Frequency’ refers to the total number of such sites across all positions considered in the Veeramah et al. *(25)* or Gronau et al. loci *(26)*.

## Supplemental Table Legends

**Table S1. General information on gibbons, whole-genome sequencing datasets and LAVA predictions.** Sex, species, and genus of 23 gibbons used to generate whole genome sequencing (WGS) libraries in this study are provided. Rows are color-coded based on genus (*Nomascus*: blue, *Hoolock*: orange, *Hylobates*: pink and *Siamang*: green). Coverage and sequencing depth are provided for each WGS dataset, along with alignment statistics. The “number of LAVA insertion loci” represents the total insertion sites in each genome, regardless of genotype at each locus. The “number of LAVA insertions in diploid genome” represents a tally of LAVA elements in each genome and across all insertion loci, considering genotypes at each locus. Thus, at each given LAVA insertion locus, homozygous LAVA insertions are counted as two inserts, and a heterozygous insertion is counted as one copy in a diploid genome.

**Table S2. LAVA insertion information and LAVA genotypes of individuals.** For each LAVA insertion, various information is reported, including the position of its insertion site, insertion direction of (−/+ for intergenic insertions and sense/antisense for genic insertions), nearest gene within 3 Kb and binary genotypes (homozygous or heterozygous presence of LAVA=1, homozygous absence of LAVA=0, unknown=NA). The *All-LAVA-all.genera* tab lists all 5,490 LAVA identified across the four genera and surviving filtrations, the *Fixed-LAVA-NLE* tab lists the 1,095 fixed LAVA insertions identified among NLE gibbons, and the *Poly-LAVA-NLE* tab includes the 1,171 poly-LAVA among NLE gibbons.

**Table S3. Transcription factor (TF) motifs significantly overrepresented in LAVA sequences.** All significantly overrepresented TF motifs (Benjamini-Hochberg corrected p-value <0.05) from TRAP and HOMER pipelines are listed in two separate tabs.

**Table S4. Histone and PU.1 ChIP-seq datasets.** General statistics for each histone and PU.1 ChIP-seq dataset (e.g. read length and count) are provided.

**Table S5. Evolutionary analysis of fixed and polymorphic LAVA.** Results from Tajima’s D analysis in windows upstream and downstream of fixed and polymorphic LAVA are presented. Comparisons between two neutral models (Supplemental Text) are presented in the model comparisons spreadsheet tab.

**Table S6. Gene ontology (GO) terms enriched near fixed LAVA**. Significant biological functions and cellular component GO terms (q<0.1) are listed along with the corresponding genes and fixed LAVA elements.

## References

1. M. K. Konkel, M. A. Batzer, A mobile threat to genome stability: The impact of non-LTR retrotransposons upon the human genome. Semin. Cancer Biol. 20, 211–221 (2010).

2. L. Schrader, J. Schmitz, The impact of transposable elements in adaptive evolution. Molecular Ecology. 28, 1537–1549 (2019).

3. G. Bourque, Transposable elements in gene regulation and in the evolution of vertebrate genomes. Current Opinion in Genetics & Development. 19, 607–612 (2009).

4. M. Trizzino, A. Kapusta, C. D. Brown, Transposable elements generate regulatory novelty in a tissue-specific fashion. BMC Genomics. 19, 468 (2018).

5. V. Sundaram, et al., Widespread contribution of transposable elements to the innovation of gene regulatory networks. Genome Res. 24, 1963–1976 (2014).

6. I. A. Warren, et al., Evolutionary impact of transposable elements on genomic diversity and lineagespecific innovation in vertebrates. Chromosome Res. 23, 505–531 (2015).

7. M. Trizzino, et al., Transposable elements are the primary source of novelty in primate gene regulation. Genome Res. 27, 1623–1633 (2017).

8. D. Blanco-Melo, R. J. Gifford, P. D. Bieniasz, Co-option of an endogenous retrovirus envelope for host defense in hominid ancestors. eLife. 6, e22519 (2017).

9. A. F. Gombart, T. Saito, H. P. Koeffler, Exaptation of an ancient Alu short interspersed element provides a highly conserved vitamin D-mediated innate immune response in humans and primates. BMC Genomics. 10, 321 (2009).

10. C. Cunningham, A. Mootnick, Gibbons. Current Biology. 19, R543–R544 (2009).

11. L. Carbone, et al., A High-Resolution Map of Synteny Disruptions in Gibbon and Human Genomes. PLOS Genetics. 2, e223 (2006).

12. L. Carbone, et al., Gibbon genome and the fast karyotype evolution of small apes. Nature. 513, 195–201 (2014).

13. L. Carbone, et al., Centromere remodeling in Hoolock leuconedys (Hylobatidae) by a new transposable element unique to the gibbons. Genome Biol Evol. 4, 648–658 (2012).

14. T. J. Meyer, et al., The Flow of the Gibbon LAVA Element Is Facilitated by the LINE-1 Retrotransposition Machinery. Genome Biol Evol. 8, 3209–3225 (2016).

15. H. Wang, et al., SVA Elements: A Hominid-specific Retroposon Family. Journal of Molecular Biology. 354, 994–1007 (2005).

16. B. Ianc, C. Ochis, R. Persch, O. Popescu, A. Damert, Hominoid Composite Non-LTR Retrotransposons—Variety, Assembly, Evolution, and Structural Determinants of Mobilization. Mol Biol Evol. 31, 2847–2864 (2014).

17. E. B. Chuong, N. C. Elde, C. Feschotte, Regulatory activities of transposable elements: from conflicts to benefits. Nat Rev Genet. 18, 71–86 (2017).

18. E. J. Gardner, et al., The Mobile Element Locator Tool (MELT): population-scale mobile element discovery and biology. Genome Res. 27, 1916–1929 (2017).

19. P. H. Sudmant, et al., An integrated map of structural variation in 2,504 human genomes. Nature. 526, 75–81 (2015).

20. T. Hara, Y. Hirai, I. Jahan, H. Hirai, A. Koga, Tandem repeat sequences evolutionarily related to SVA-type retrotransposons are expanded in the centromere region of the western hoolock gibbon, a small ape. J. Hum. Genet. 57, 760–765 (2012).

21. Y.-C. Chan, et al., Mitochondrial Genome Sequences Effectively Reveal the Phylogeny of Hylobates Gibbons. PLOS ONE. 5, e14419 (2010).

22. K. Matsudaira, T. Ishida, Phylogenetic relationships and divergence dates of the whole mitochondrial genome sequences among three gibbon genera. Molecular Phylogenetics and Evolution. 55, 454–459 (2010).

23. K. R. Veeramah, et al., Examining phylogenetic relationships among gibbon genera using whole genome sequence data using an approximate bayesian computation approach. Genetics. 200, 295–308 (2015).

24. C.-M. Shi, Z. Yang, Coalescent-Based Analyses of Genomic Sequence Data Provide a Robust Resolution of Phylogenetic Relationships among Major Groups of Gibbons. Mol. Biol. Evol. 35, 159–179 (2018).

25. C. Gao, et al., Characterization and functional annotation of nested transposable elements in eukaryotic genomes. Genomics. 100, 222–230 (2012).

26. T. Sultana, A. Zamborlini, G. Cristofari, P. Lesage, Integration site selection by retroviruses and transposable elements in eukaryotes. Nature Reviews Genetics. 18, 292–308 (2017).

27. G. Bourque, et al., Ten things you should know about transposable elements. Genome Biology. 19, 199 (2018).

28. N. H. Lazar, et al., Epigenetic maintenance of topological domains in the highly rearranged gibbon genome. Genome Res. 28, 983–997 (2018).

29. L. Li, Z. Wunderlich, An Enhancer’s Length and Composition Are Shaped by Its Regulatory Task. Front Genet. 8(2017).

30. G. Bourque, et al., Evolution of the mammalian transcription factor binding repertoire via transposable elements. Genome Res. 18, 1752–1762 (2008).

31. P. Rimmelé, et al., Spi-1/PU.1 Oncogene Accelerates DNA Replication Fork Elongation and Promotes Genetic Instability in the Absence of DNA Breakage. Cancer Res. 70, 6757–6766 (2010).

32. S. W. Criscione, Y. Zhang, W. Thompson, J. M. Sedivy, N. Neretti, Transcriptional landscape of repetitive elements in normal and cancer human cells. BMC Genomics. 15, 583 (2014).

33. J. D. Fernandes, et al., The UCSC repeat browser allows discovery and visualization of evolutionary conflict across repeat families. Mobile DNA. 11, 13 (2020).

34. ENCODE Project Consortium, An integrated encyclopedia of DNA elements in the human genome. Nature. 489, 57–74 (2012).

35. C. A. Davis, et al., The Encyclopedia of DNA elements (ENCODE): data portal update. Nucleic Acids Res. 46, D794–D801 (2018).

36. I. Lupan, P. Bulzu, O. Popescu, A. Damert, Lineage specific evolution of the VNTR composite retrotransposon central domain and its role in retrotransposition of gibbon LAVA elements. BMC Genomics. 16, 389 (2015).

37. O. Gianfrancesco, V. J. Bubb, J. P. Quinn, SVA retrotransposons as potential modulators of neuropeptide gene expression. Neuropeptides. 64, 3–7 (2017).

38. D. J. Elzi, M. Song, K. Hakala, S. T. Weintraub, Y. Shiio, Wnt Antagonist SFRP1 Functions as a Secreted Mediator of Senescence. Molecular and Cellular Biology. 32, 4388–4399 (2012).

39. Z. Zhou, J. Wang, X. Han, J. Zhou, S. Linder, Up-regulation of human secreted frizzled homolog in apoptosis and its down-regulation in breast tumors. Int. J. Cancer. 78, 95–99 (1998).

40. F. Tajima, Statistical method for testing the neutral mutation hypothesis by DNA polymorphism. Genetics. 123, 585–595 (1989).

41. R. N. Gutenkunst, R. D. Hernandez-Rodriguez, S. H. Williamson, C. D. Bustamante, Inferring the Joint Demographic History of Multiple Populations from Multidimensional SNP Frequency Data.

42. K. Thornton, Recombination and the Properties of Tajima’s D in the Context of Approximate-Likelihood Calculation. Genetics. 171, 2143–2148 (2005).

43. S. X. Pfister, et al., SETD2-Dependent Histone H3K36 Trimethylation Is Required for Homologous Recombination Repair and Genome Stability. Cell Rep. 7, 2006–2018 (2014).

44. F.-L. Tsai, M. Kai, The checkpoint clamp protein Rad9 facilitates DNA-end resection and prevents alternative non-homologous end joining. Cell Cycle. 13, 3460–3464 (2014).

45. R. K. Pandita, et al., Mammalian Rad9 Plays a Role in Telomere Stability, S- and G2-Phase-Specific Cell Survival, and Homologous Recombinational Repair. Mol Cell Biol. 26, 1850–1864 (2006).

46. J. An, E. González-Avalos, et al., Acute loss of TET function results in aggressive myeloid cancer in mice. Nature Communications. 6, 1–14 (2015).

47. D. Jiang, S. Wei, F. Chen, Y. Zhang, J. Li, TET3-mediated DNA oxidation promotes ATR-dependent DNA damage response. EMBO Rep. 18, 781–796 (2017).

48. T. Rampias, et al., The lysine-specific methyltransferase KMT2C/MLL3 regulates DNA repair components in cancer. EMBO Rep. 20(2019).

49. M. Y. Shah, et al., MMSET/WHSC1 enhances DNA damage repair leading to an increase in resistance to chemotherapeutic agents. Oncogene. 35, 5905–5915 (2016).

50. K. H. Kim, C. W. M. Roberts, Mechanisms by which SMARCB1 loss drives rhabdoid tumor growth. Cancer Genet. 207, 365–372 (2014).

51. J. O. Yáñez-Cuna, E. Z. Kvon, A. Stark, Deciphering the transcriptional cis-regulatory code. Trends in Genetics. 29, 11–22 (2013).

52. M. Ridinger-Saison, et al., Spi-1/PU.1 activates transcription through clustered DNA occupancy in erythroleukemia. Nucleic Acids Res. 40, 8927–8941 (2012).

53. D. Ezer, N. R. Zabet, B. Adryan, Homotypic clusters of transcription factor binding sites: A model system for understanding the physical mechanics of gene expression. Computational and Structural Biotechnology Journal. 10, 63–69 (2014).

54. A. D. Ewing, Transposable element detection from whole genome sequence data. Mob DNA. 6 (2015)

55. S. Carvalho, et al., SETD2 is required for DNA double-strand break repair and activation of the p53-mediated checkpoint. eLife. 3, e02482 (2014).

56. P. Goerner-Potvin, G. Bourque, Computational tools to unmask transposable elements. Nature Reviews Genetics. 19, 688 (2018).

57. H. Li, R. Durbin, Fast and accurate short read alignment with Burrows-Wheeler transform. Bioinformatics. 25, 1754–1760 (2009).

58. S. Andrews, FastQC: A quality control tool for high throughput sequence data. (2010) (available at http://www.bioinformatics.babraham.ac.uk/projects/fastqc/).

59. F. Ramírez, F. Dündar, S. Diehl, B. A. Grüning, T. Manke, deepTools: a flexible platform for exploring deep-sequencing data. Nucleic Acids Res. 42, W187–W191 (2014).

60. J. Ernst, M. Kellis, Chromatin-state discovery and genome annotation with ChromHMM. Nature Protocols. 12, 2478–2492 (2017).

61. A. R. Quinlan, I. M. Hall, BEDTools: a flexible suite of utilities for comparing genomic features. Bioinformatics. 26, 841–842 (2010).

62. S. Heinz, et al., Simple Combinations of Lineage-Determining Transcription Factors Prime cis-Regulatory Elements Required for Macrophage and B Cell Identities. Molecular Cell. 38, 576–589 (2010).

63. M. Thomas-Chollier, et al., Transcription factor binding predictions using TRAP for the analysis of ChIP-seq data and regulatory SNPs. Nature Protocols. 6, 1860–1869 (2011).

64. Y. H. Y Benjamini, Controlling the false fiscovery rate: a practical and powerful approach to multiple testing. J. Royal Statist. Soc., Series B. 57, 289–300 (1995).

65. A. A. Shabalin, Matrix eQTL: ultra fast eQTL analysis via large matrix operations. Bioinformatics. 28, 1353–1358 (2012).

66. T. S. Korneliussen, A. Albrechtsen, R. Nielsen, ANGSD: Analysis of Next Generation Sequencing Data. BMC Bioinformatics. 15, 356 (2014).

67. 1000 Genomes Project Consortium, A global reference for human genetic variation. Nature. 526, 68–74 (2015).

68. M. V. Kuleshov, et al., Enrichr: a comprehensive gene set enrichment analysis web server 2016 update. Nucleic Acids Res. 44, W90–W97 (2016).

## Supplemental References

1. E. J. Gardner, et al., The Mobile Element Locator Tool (MELT): population-scale mobile element discovery and biology. Genome Res. 27, 1916–1929 (2017).

2. A. R. Quinlan, I. M. Hall, BEDTools: a flexible suite of utilities for comparing genomic features. Bioinformatics. 26, 841–842 (2010).

3. H. Li, R. Durbin, Fast and accurate short read alignment with Burrows-Wheeler transform. Bioinformatics. 25, 1754–1760 (2009).

4. H. Li, et al., The Sequence Alignment/Map format and SAMtools. Bioinformatics. 25, 2078–2079 (2009).

5. D. Kostka, A. K. Holloway, K. S. Pollard, Developmental Loci Harbor Clusters of Accelerated Regions That Evolved Independently in Ape Lineages. Mol Biol Evol. 35, 2034–2045 (2018).

6. V. J. Lynch et al. Ancient transposable elements transformed the uterine regulatory landscape and transcriptome during the evolution of mammalian pregnancy. Cell Rep. 10, 551–561 (2015).

7. Y. H. Y Benjamini, Controlling the false fiscovery rate: a practical and powerful approach to multiple testing. J. Royal Statist. Soc., Series B. 57, 289–300 (1995).

8. S. Neph, et al. Stamatoyannopoulos, BEDOPS: high-performance genomic feature operations. Bioinformatics. 28, 1919–1920 (2012).

9. L. Carbone, et al., Gibbon genome and the fast karyotype evolution of small apes. Nature. 513, 195–201 (2014).

10. N. H. Lazar, et al., Epigenetic maintenance of topological domains in the highly rearranged gibbon genome. Genome Res. 28, 983–997 (2018).

11. S. Andrews, FastQC: A quality control tool for high throughput sequence data. (2010) (available at http://www.bioinformatics.babraham.ac.uk/projects/fastqc/).

12. A. M. Bolger, M. Lohse, B. Usadel, Trimmomatic: a flexible trimmer for Illumina sequence data. Bioinformatics. 30, 2114–2120 (2014).

13. S. W. Criscione, Y. Zhang, W. Thompson, J. M. Sedivy, N. Neretti, Transcriptional landscape of repetitive elements in normal and cancer human cells. BMC Genomics. 15, 583 (2014).

14. M. D. Robinson, D. J. McCarthy, G. K. Smyth, edgeR: a Bioconductor package for differential expression analysis of digital gene expression data. Bioinformatics. 26, 139–140 (2010).

15. J. D. Fernandes, et al. The UCSC repeat browser allows discovery and visualization of evolutionary conflict across repeat families. Mobile DNA. 11, 13 (2020).

16. R. C. Edgar, MUSCLE: multiple sequence alignment with high accuracy and high throughput. Nucleic Acids Res. 32, 1792–1797 (2004).

17. G. Benson, Tandem repeats finder: a program to analyze DNA sequences. Nucleic Acids Res. 27, 573–580 (1999).

18. W. J. Kent, BLAT--the BLAST-like alignment tool. Genome Res. 12, 656–664 (2002).

19. Y. Zhang, et al., Model-based Analysis of ChIP-Seq (MACS). Genome Biology. 9, R137 (2008).

20. F. Ramírez, F. Dündar, S. Diehl, B. A. Grüning, T. Manke, deepTools: a flexible platform for exploring deep-sequencing data. Nucleic Acids Res. 42, W187–W191 (2014).

21. A. Dobin, et al., STAR: ultrafast universal RNA-seq aligner. Bioinformatics. 29, 15–21 (2013).

22. K. D. Hansen, R. A. Irizarry, Z. Wu, Removing technical variability in RNA-seq data using conditional quantile normalization. Biostatistics. 13, 204–216 (2012).

23. M. I. Love, W. Huber, S. Anders, Moderated estimation of fold change and dispersion for RNA-seq data with DESeq2. Genome Biol. 15, 550 (2014).

24. T. S. Korneliussen, A. Albrechtsen, R. Nielsen, ANGSD: Analysis of Next Generation Sequencing Data. BMC Bioinformatics. 15, 356 (2014).

25. K. R. Veeramah, et al., Examining phylogenetic relationships among gibbon genera using whole genome sequence data using an approximate bayesian computation approach. Genetics. 200, 295–308 (2015).

26. I. Gronau, M. J. Hubisz, B. Gulko, C. G. Danko, A. Siepel, Bayesian inference of ancient human demography from individual genome sequences. Nature Genetics. 43, 1031–1034 (2011).

27. F. Tajima, Statistical method for testing the neutral mutation hypothesis by DNA polymorphism. Genetics. 123, 585–595 (1989).

28. R. N. Gutenkunst, R. D. Hernandez-Rodriguez, S. H. Williamson, C. D. Bustamante, Inferring the Joint Demographic History of Multiple Populations from Multidimensional SNP Frequency Data.

29. K. Thornton, Recombination and the Properties of Tajima’s D in the Context of Approximate-Likelihood Calculation. Genetics. 171, 2143–2148 (2005).

30. 1000 Genomes Project Consortium, A global reference for human genetic variation. Nature. 526, 68–74 (2015).

31. M. V. Kuleshov, et al., Enrichr: a comprehensive gene set enrichment analysis web server 2016 update. Nucleic Acids Res. 44, W90–W97 (2016).

